# The Role of Metabolism in Shaping Enzyme Structures Over 400 Million Years of Evolution

**DOI:** 10.1101/2024.05.27.596037

**Authors:** Oliver Lemke, Benjamin Murray Heineike, Sandra Viknander, Nir Cohen, Jacob Lucas Steenwyk, Leonard Spranger, Feiran Li, Federica Agostini, Cory Thomas Lee, Simran Kaur Aulakh, Jens Nielsen, Antonis Rokas, Judith Berman, Aleksej Zelezniak, Toni Ingolf Gossmann, Markus Ralser

## Abstract

The functions of cells and proteins depend on their biochemical microenvironment. To understand how biochemical constraints shaped protein structural evolution, we coupled the extensive genetic and metabolic data from the *Saccharomycotina* subphylum with the capability of AlphaFold2 to systematically predict protein structures from sequence. Determining how 11,269 enzyme structures catalysing 361 different metabolic reactions evolved over 400 million years alongside their molecular functions, we report that metabolism has shaped the structural evolution of enzymes at different levels: the organism’s overall metabolism; the topological organisation of the metabolic network; and each enzyme’s molecular properties. For example, structural evolution depends on each enzyme’s reaction mechanism, on the variability rather than the amount of metabolic flux, and on biosynthetic cost. Evolutionary cost-optimization is stronger on highly abundant enzymes and acts differently on different structural domains, with the exception of small-molecule binding sites, which are prioritised over other structural domains and lack cost-optimisation. Finally, while enzyme surfaces are less constrained, surface residues can also be exposed to positive selection for the co-evolution of protein-protein interaction sites. Accessing AlphaFold’s power to predict protein structures systematically and across species barriers, facilitating the integration of protein structures with functional genomics, we were thus able to map biological constraints which shape protein structural evolution at scale and over long timelines.

## Introduction

Metabolism drives biological energy conversion, and determines the metabolite composition of cells, constraining how they evolve, function, and withstand stressful conditions. Cells adjust the activity of their enzymes to fit their environment and thus optimise their use of available nutrients and adjust to external perturbations in their niche (Afriat et al., 2006; Voordeckers et al., 2012; Yang et al., 2019; Yoshida et al., 2016). Enzyme plasticity is important for across disciplines, including the bioengineering of novel or improved metabolic reactions (Porter et al., 2016). In medicine, enzyme plasticity applies to drug responses in inherited and acquired metabolic disease, cancer and microbial infections (Lopatkin et al., 2021; Pavlova and Thompson, 2016; Pérez et al., 2012). In drug discovery, enzymes represent a major class of drug targets (Stine et al., 2022). Despite the undisputed importance of enzyme dynamics, we have an incomplete picture of the biochemical constraints acting on their function and evolution.

Enzymes function as part of the metabolic network, a large, interconnected biological system that has evolved both at the network topology and individual enzyme levels (Guzmán et al., 2019; Noda-Garcia et al., 2018; Ralser et al., 2021). With our increasing knowledge of the former, it is presumed that the metabolic network structure has been shaped by biophysical and thermodynamic parameters, that include optimality principles (Ng et al., 2019), as well as enzyme interactions with cellular metabolites, which often act as inhibitors or activators (Alam et al., 2017; Zubaidi et al., 2023). Furthermore, the intrinsic chemical reaction principles of the interconverted metabolites shaped the evolution of metabolic pathways since their origin. For example, many enzyme-catalysed reactions of central metabolic pathways, including glycolysis, the tricarboxylic acid cycle, and the pentose phosphate pathway, have non-enzymatic counterparts that are promoted by metal ions, particularly Fe(II), which was abundant in Archean sediment (Keller et al., 2016; Muchowska et al., 2020; Rouxel et al., 2005). Thus, a likely scenario is that many reactions that now participate in the central metabolic network originated on the basis of non-enzymatic, often metal-catalysed reactions, that accelerated and became regulated by the evolution of protein-based enzymes in early organisms (Aulakh et al., 2022; Keller et al., 2016; Muchowska et al., 2020; Ralser, 2018). As a consequence, enzymes evolve alongside a topological network organisation that stems largely from the chemical reaction properties of the interconverted metabolites.

For species that existed after the last universal common ancestor, ancestral metabolic networks and enzymes can be reconstructed on the basis of sequence comparisons (Clifton et al., 2018; Mascotti, 2022; Nicoll et al., 2020; Voordeckers et al., 2012). At the network level, these approaches can provide information on the metabolic properties or habitats of ancestral species, and at the enzyme level, information on the evolution of substrate specificities can be obtained (Goldford et al., 2017; Goldford and Segrè, 2018; Weiss et al., 2016). More recent evolutionary events can be reconstructed in greater detail. For instance, species-specific metabolic models in *Saccharomycotina* yeasts have revealed how metabolic capabilities are gained within clades, often through the expansion of existing gene families and the increased substrate specificity of promiscuous enzymes (Lu et al., 2021).

Also, the evolution of enzyme structures themselves has thus far been studied mostly through the lens of comparative genomics. Comparing enzyme sequences across evolution has revealed the impact of various constraints at the amino acid level, such as preserving the chemical identity of amino acid side chains and epistatic interactions between residues (Breen et al., 2012; Suárez-Díaz, 2016; Zuckerkandl and Pauling, 1965). Other studies that integrated metabolic network reconstructions revealed how the substantial costs enzyme biosynthesis imposes on the biological system are a major constraint in protein evolution (Chen and Nielsen, 2022). Biological cost optimization is observed at the species and molecular levels. For example, in many conditions cells prefer cost-effective fermentation over oxidative metabolism for the production of ATP, despite the latter producing stoichiometrically higher ATP amounts (Nilsson and Nielsen, 2016). Moreover, many enzymes are amongst the most abundant proteins, and have evolved by incorporating less energetically costly amino acids (Akashi and Gojobori, 2002; Barton et al., 2010; Raiford et al., 2008).

We speculated that protein structures could augment our understanding of these relationships, by providing information on the co-localization of residues within proteins, distinguish surface from core residues, and enable the identification of functional sites, such as active sites, binding pockets for allosteric regulators, and protein-protein interaction sites (Studer et al., 2013) that might be exposed to specific selective constraints. In order to identify the metabolic constraints affecting enzyme structural evolution, we used a combination of comparative evolutionary genomic and comparative structural biology, exploiting the extensive metabolic and genetic characterisation of the *Saccharomycotina* subphylum, which contains the biotechnologically and medically relevant yeasts *Saccharomyces cerevisiae* and *Candida albicans* (Kurtzman et al., 2011; Lu et al., 2021; Opulente et al., 2024; Shen et al., 2018), as well as recent advances in computational structural biology that allow us to predict high quality protein structure at scale and across species barriers using Alphafold2 (Jumper et al., 2021). To test how the divergent metabolic properties between these species, their metabolic pathways, and the different enzyme reaction mechanisms represented in the metabolic network acted on protein structural evolution, we used AlphaFold2 to assemble structures of 11,269 enzymes from a selection of 27 species, which evolved over 400 million years (Opulente et al., 2024; Shen et al., 2018). These are clustered into 424 groups of orthologs (orthogroups) covering 361 different metabolic reactions as represented by different Enzyme Commission (EC) classifiers, participating in 224 metabolic pathways. We found that the properties of the underlying metabolic networks have influenced enzyme structural evolution at different scales, ranging from the level of the organism and its overall metabolic niche to the level of metabolic pathways and catalysed reactions, and even the molecular level.

## Results

### Mapping evolutionary adaptations in enzyme structures over 400 million years of evolution

Out of a panel of 332 genome-sequenced species belonging to the *Saccharomycotina* subphylum (Shen et al., 2018; Wu et al., 2017) we selected 26 highly phylogenetically diverse genomes (Supplementary Note 1). We further included the model fission yeast *Schizosaccharomyces pombe* as an outgroup to help root protein trees (Figure 1A, Table S1.1). We then selected the sequences of metabolic enzymes, based on sequence homology with metabolic enzymes in the Yeast Pathways Database (“YeastPathways Database Website Home,” 2023), using the OrthoMCL clusters generated by Shen et al. (2018). This resulted in 424 orthogroups, for which we predicted or assembled Alphafold2-structures comprising 11,269 enzymes in total (n = 1,301 structures we extracted from AlphafoldDB (Varadi et al., 2022) and n = 9,968 we predicted at the Berzelius computing infrastructure, using AlphaFold version 2.0.1 at the start of our project, Supplementary Note 2). From our original set of orthologs (n = 10,545 sequences) we were unable to predict structures for 577 enzyme sequences. 59.3% of the unpredicted sequences were from just three species, *Hanseniaspora osmophelia*, *Zygosaccharomyces rouxii*, and *Yarrowia lipolytica*. The unpredicted structures also denote longer sequences (Table S1.2), and no structures were predicted for two orthogroups, including the *S. cerevisiae* acetyl-coenzyme A carboxylases Hfa1p and Acc1p, as well as 3-hydroxyacyl-CoA dehydrogenase and enoyl-CoA hydratase Fox2p (Table S1.3). The final predicted structures covered 44 % (361/817) of all EC classifiers, distributed over 224 metabolic pathways in *S. cerevisiae* (Table S1.4). This selection encompassed all major enzyme classes, including 119 oxidoreductases, 191 transferases, 55 hydrolases, 49 lyases, 21 isomerases, 29 ligases, and 2 translocases.

**Figure 1:**
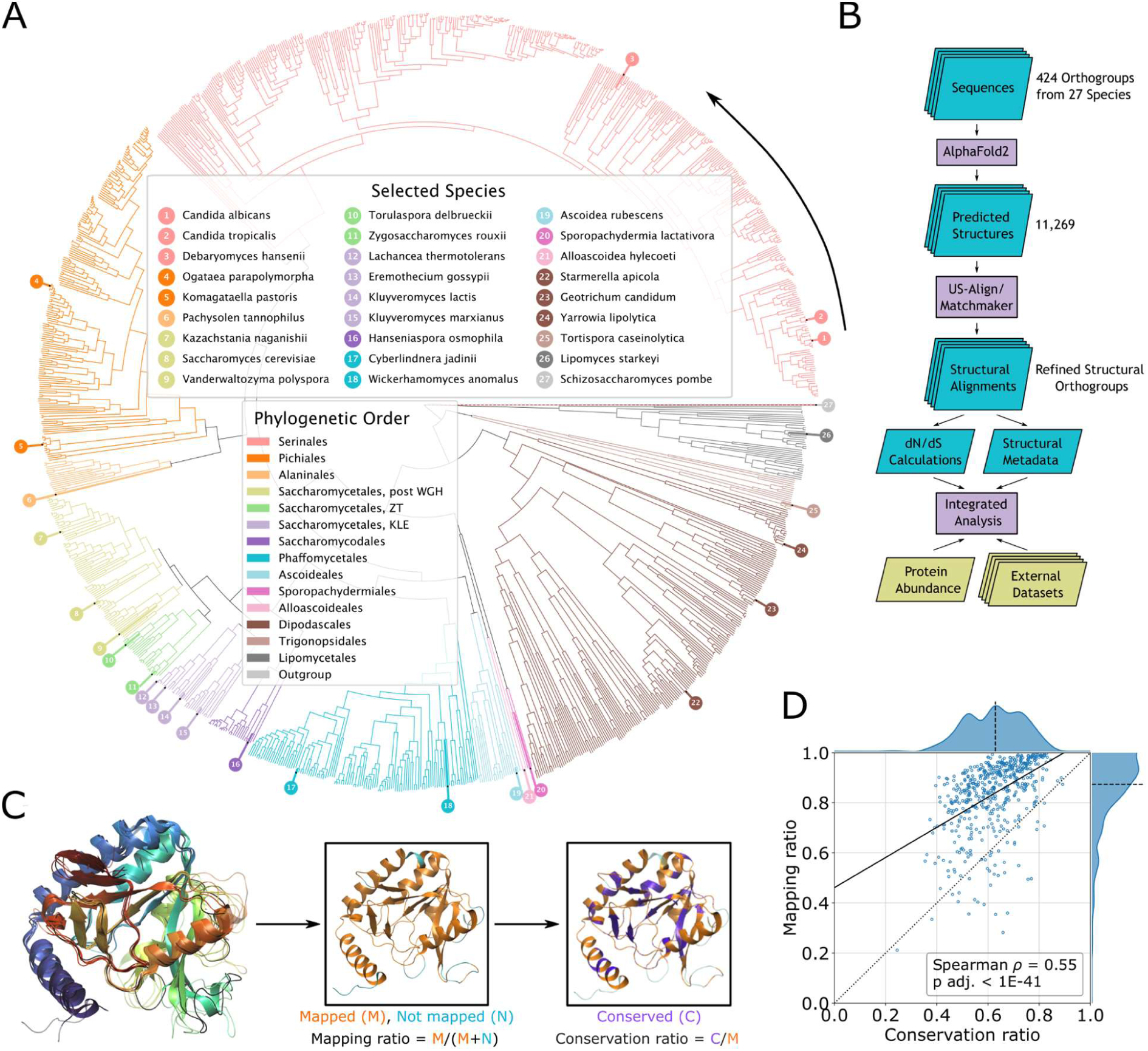
Comparing Alphafold2-predicted structures of 11,269 metabolic enzymes from 27 species of the *Saccaromycotina spp*. to metabolic properties at species, pathway and molecular levels reveal properties of protein structural evolution. A) Phylogenetic tree of the *Saccharomycotina* yeast subphylum highlighting the 27 species from which metabolic enzyme sequences were extracted and used for structural prediction, coloured as indicated by phylogenetic order, numbering counterclockwise starting at *C. albicans*. Branch lengths and topology are from the species time tree as calculated in (Opulente et al., 2024), except for the branch for the outgroup species, *S. pombe* which is not drawn to scale. B) Illustration of our analysis pipeline. C) Example alignment for 5-Formyltetrahydrofolate cyclo-ligase (Fau1p in *S. cerevisiae*) in 5 species (*S. cerevisiae*, *C. albicans*, *K. lactis*, *K. pastoris* and *S. pombe*). The black line denotes the reference structure in *S. cerevisiae*. Insets show the ortholog from *C. albicans* with mapped residues (orange), unmapped residues (cyan) as well as residues conserved between *S. cerevisiae* and *C. albicans* (purple). D) Mean mapping ratio versus mean conservation ratio with respect to 529 reference structures that passed our filters. Dotted line denotes the identity line, the dashed line denotes the axis median and the solid line indicates the best linear fit.

We assessed the prediction quality using the predicted Local Distance Difference Test (pLDDT) score, a per-residue confidence estimate provided by AlphaFold2, for all enzymes (range: [0,100]; best: 100) (Guo et al., 2022; Mariani et al., 2013). Overall, our dataset included well-structured protein structures with a mean pLDDT of 90.4 and mean coefficient of variation (CV) of 0.15, which was mostly affected by terminal regions (first 10% of the sequence: mean pLDDT = 79.1, mean CV = 0.29; central 80%: mean pLDDT = 92.2, mean CV = 0.13; last 10%: mean pLDDT = 87.3, mean CV = 0.18) (Figure S1.1A). To validate the sequence-based orthogroup assignments generated by Shen et al., (2018), we used hierarchical clustering based on a bidirectional template modelling score (TM-score) (Zhang et al., 2022; Zhang and Skolnick, 2005) (Figures S1.1B,C). Indeed, 29 sequence-based orthogroups were split into distinct structural orthogroups (Table S1.5).

To better benefit from the vast amount of metabolic and phenotypic data available for the model yeast *S. cerevisiae,* we also calculated separate pairwise alignments to *S. cerevisiae* enzyme structures within each reclustered structural orthogroup using the Matchmaker algorithm in Chimera (Meng et al., 2006). From these pairwise structural alignments, we obtained metadata including a conservation ratio (CR), mapping ratio (MR), amino acid constitution, and other structure-specific properties such as solvent-accessible surface area (SASA), relative location within the protein, and secondary structure composition (Table S1.6). We analysed the obtained metadata together with calculated dN/dS ratios and data from external resources, including public databases, known crystal structures, and experimental datasets to link the structural properties to the phenotypic and metabolic properties for each orthogroup (Figure 1B; see Table 1 in Methods).

**Table 1:**
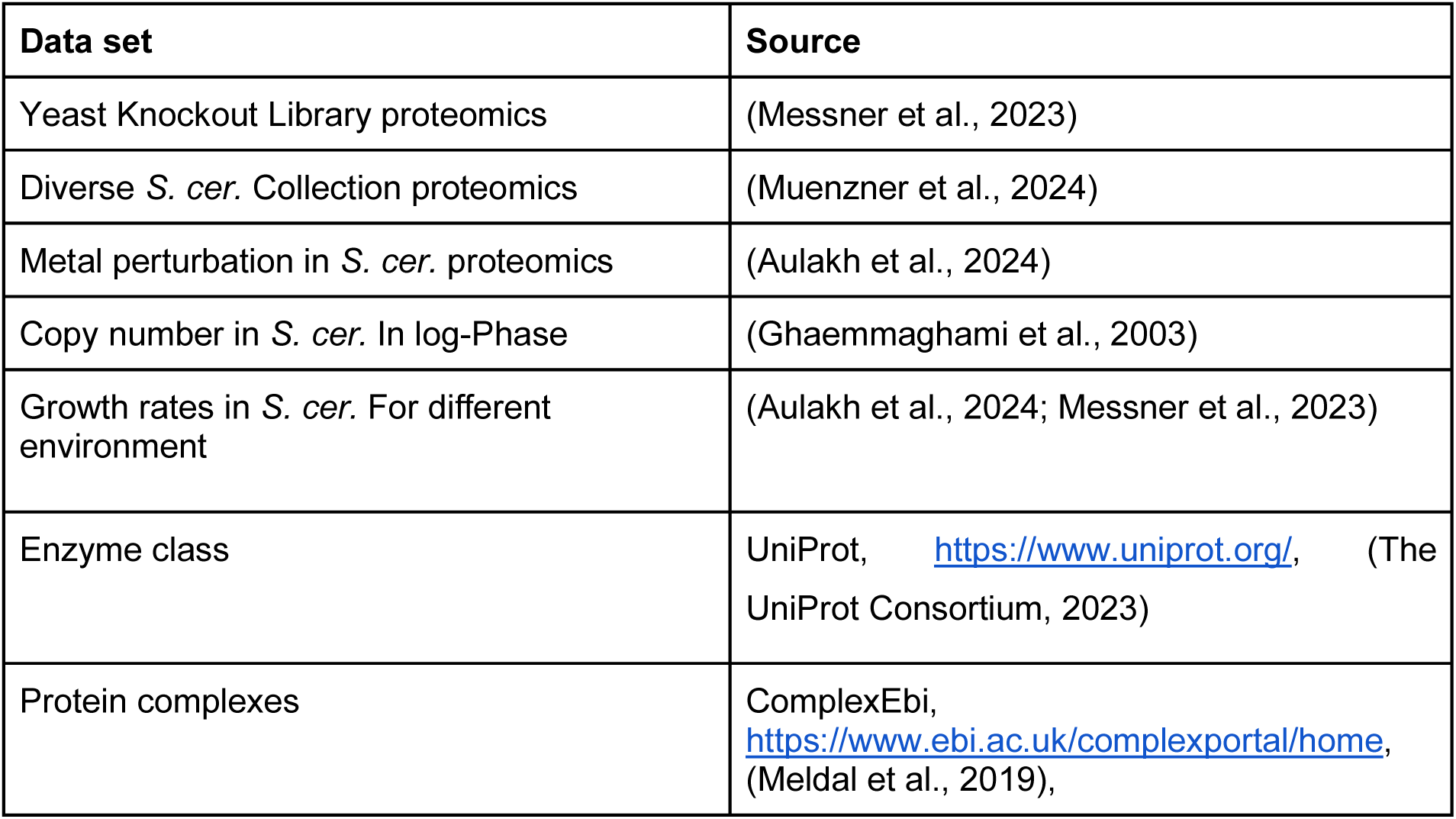
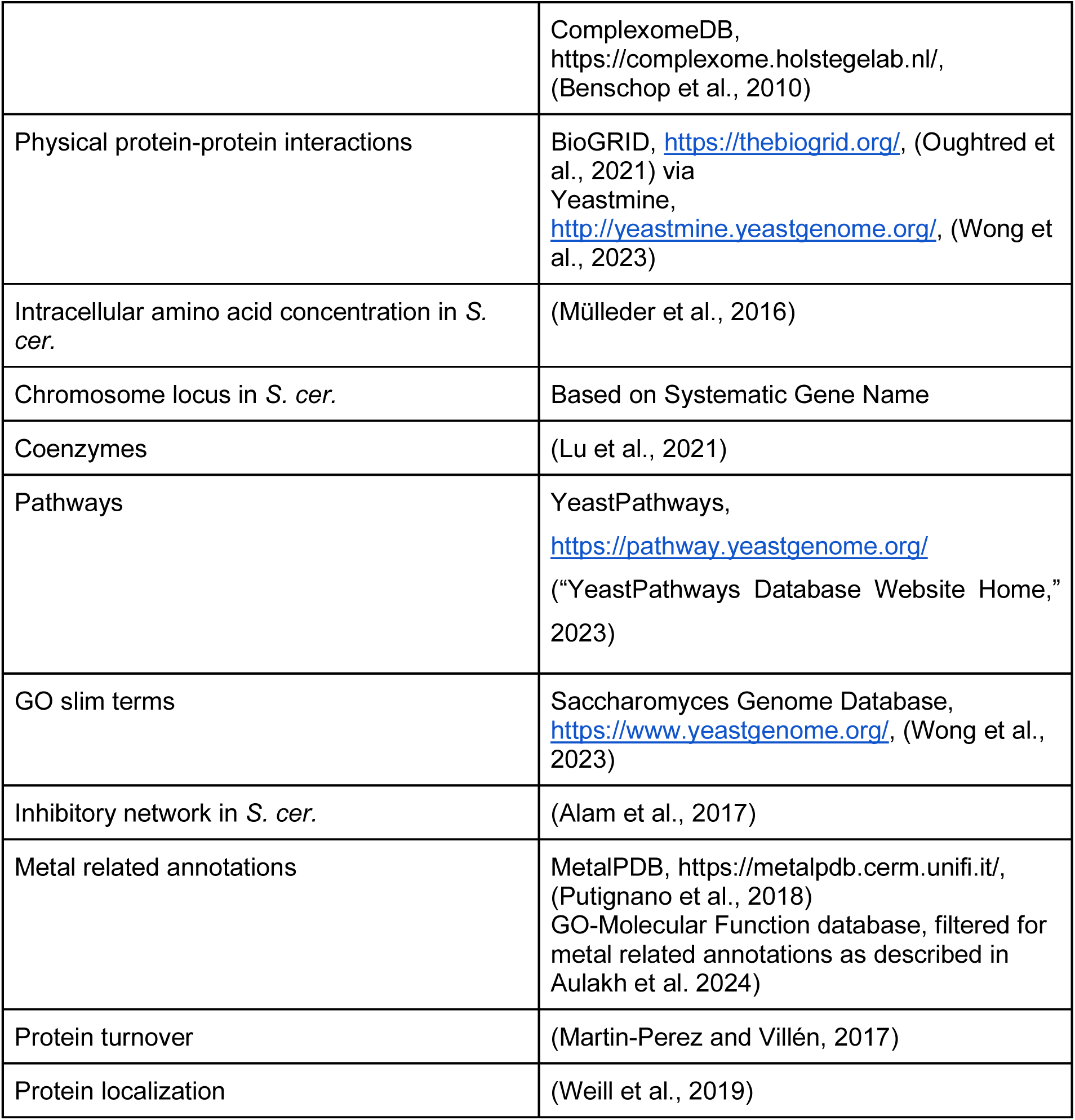
Datasets used throughout the presented research.

To determine the degree of enzyme structural diversification among the 27 species representing the subphylum, we calculated an averaged MR and an averaged CR for each orthogroup using designated *S. cerevisiae* reference structures for comparison. The MR quantifies the percentage of amino acids that can be 1:1 mapped to a structure of *S. cerevisiae* enzyme. The CR quantifies the percentage of mapped residues that are identical to those of the reference structure. Both metrics are illustrated with the orthogroup of 5-Formyltetrahydrofolate cyclo-ligase (27 structures, CR = 0.40, MR = 0.87, Figures 1C and S1.1D, Fau1p in *S. cerevisiae*), which is involved in tetrahydrofolate metabolism (Holmes and Appling, 2002). In this orthogroup, the secondary structural elements (helical, extended) showed a mean MR of 95.4% whereas the longer random coil regions, which have a higher conformational flexibility, only showed a mean MR of 77.3%. The median MR for all orthogroups was 87.4% (inter quartile range [IQR]: [78.3%, 93.9%]) (Figure 1D) mostly affected by low-pLDDT scoring regions (Figure S1.1E), i.e. the termini (especially the N-terminus, Figure S1.1A) and longer random coil regions (60 % of the unmapped vs. 36% of the mapped regions), which could be grouped neither to helical nor extended structures. As expected, within the orthogroups, the core structures of the enzymes overlapped, allowing them to be aligned. MR correlated with CR (Spearman ⍴ = 0.55, adj. p-value < 1E-41) (Figure 1D), which shows a median CR of 62.9% (IQR: [53.6%, 71.2%], total range: [24.5% to 89.2%]) indicating a high degree of structural diversity that could be linked to metabolic evolution.

### Biochemical constraints of enzyme structural evolution

We started our analysis at the species level and asked whether metabolic specialisations have left a signature on the protein structures. We first subgrouped the orthogroups according to the growth properties of the yeast species on 21 different carbon sources (Kurtzman et al., 2011; Lu et al., 2021; Opulente et al., 2018; Shen et al., 2018). Enzyme structures diversified depending on the preferred carbon source of the species. Enzymes in species that can grow anaerobically on glucose (Figure S2.1A), raffinose, galactose and sucrose exhibited the most significant structural diversity between subgroups, alongside enzymes from species that grow aerobically on xylose (adj. p-values < 1e-83, Wilcoxon signed-rank test). Enzyme structures from anaerobically fermenting species showed higher conservation relative to the structure from *S. cerevisiae,* which also ferments those sugars, than enzyme structures from species that prefer oxidative metabolism (Figure 2A). Indeed, for glucose-fermenting species, some of the largest differences in structural divergence between subgroups were detected within the orthogroups of enzymes implicated in central carbon metabolism or the electron transport chain (ETC). For example, Kgd2p of the TCA cycle and Cox7p of the respiratory chain, were among the enzymes exhibiting highest change in structural diversity between the fermenting and non-fermenting species. These more divergent orthogroups also included Met10p of the methionine and sulphur cycle; Ath1p of trehalose metabolism, which in yeast serves as carbon storage; and Erg1p, involved in the biosynthesis of fungi’s functional equivalent of cholesterol, ergosterol, a main drug target in the development of antifungals (Forastiero et al., 2013; Leber et al., 2003). We also observed a few exceptions; in non-fermenting species, a higher CR was detected in the orthogroups of Ndi1p, an NADH dehydrogenase of mitochondria which is a functional analogue of the respiratory chain complex I in other organisms; the aldehyde dehydrogenase Ald5p; the isocitrate dehydrogenase Idp1; and Ilv6p, which is involved in branched-chain amino acid metabolism.

**Figure 2:**
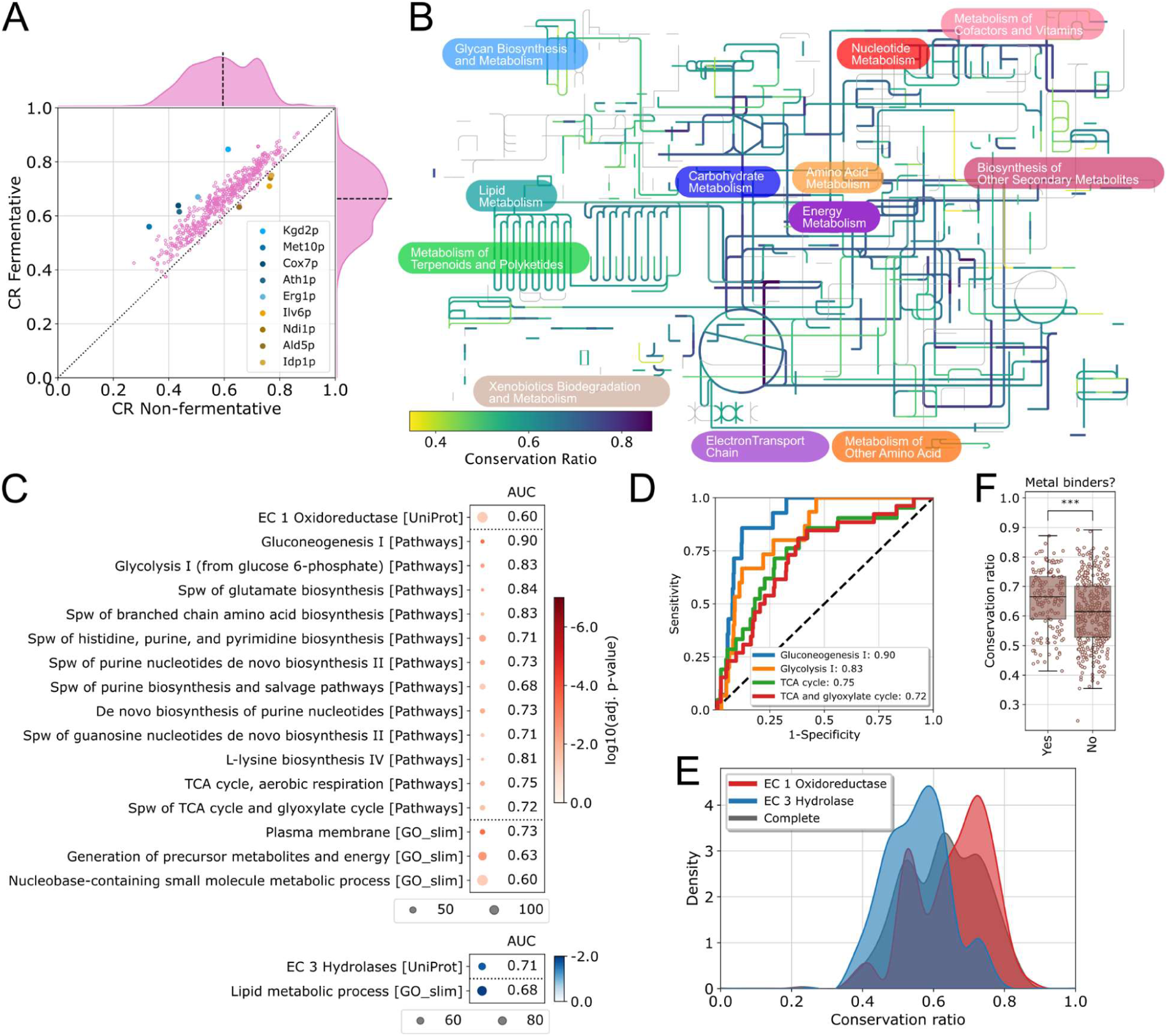
Biochemical constraints of enzyme structural evolution in *Saccharomycotina.* A) Mean conservation ratio per protein calculated for species that ferment glucose versus species that prefer an oxidative metabolism. Proteins with the largest differences in both directions are highlighted. The dotted line denotes the identity line and the dashed line denotes the axis median. B) Conservation ratio projected onto the yeast metabolic map of iPath3 (Darzi et al., 2018). C) Enrichment of the orthogroups containing the 25% structurally most conserved (red, upper panel) and the 25% structurally most diverse enzymes (blue, lower panel) for metabolic pathways from the yeast pathways database (“YeastPathways Database Website Home,” 2023) and GO slim terms. Size indicates the number of genes associated with the term or pathway in our selection, and colour indicates adjusted p-value. In addition, the area under the receiver operating characteristic (ROC) curve (abbrev. AUC) is shown. Super pathway is abbreviated as “Spw” D) ROC-curve for four highly enriched pathways covering TCA cycle and glucose metabolism. The number in the legend indicates the AUC. The dashed line denotes the identity line representing random sampling. E) Distribution of the mean conservation ratio of orthogroups assigned to oxidoreductases (red) or hydrolases (blue). The average distribution for all orthogroups is shown in black. F) Boxplot of the conservation ratio of enzymes known to bind metals and those not known to bind metals. The boxes denote the first and third quartile, the whiskers extend up to 1.5-times the interquartile range. Each dot represents an orthogroup. *** indicates a p-value < 1e-4, Wilcoxon-Mann-Whitney test.

In parallel, we performed GO-slim term enrichment analysis of the first quartile of the orthogroups with the largest differences in CR between subgroups for glucose fermentation. We obtained an enrichment for the terms “membrane”, “lipid metabolism”, “endoplasmic reticulum” and “endomembrane system” (adj. p < 1e-2, Fisher’s exact test) (Figure S2.1B). We also tested for significant CR differences between subgroups at the pathway level. We found that aerobic respiration, glucose fermentation, purine biosynthesis and ergosterol biosynthesis differed significantly between fermenting and aerobically growing species (adj. p < 1e-4, Wilcoxon signed-rank test). Thus, species that specialise metabolically showed different patterns of enzyme structural divergence. Therefore, we concluded that the metabolic niche specialisation influences the structural evolution of its enzymes.

Next, we focused on the metabolic constraints acting at the pathway level. We projected the degree of structural divergence calculated using all species within each orthogroup, onto a genome-scale reconstruction of the metabolic network (Figure 2B). Then, we performed a pathway enrichment analysis, considering the orthogroups containing the 25% most structurally divergent enzymes, as well as the 25% most conserved enzyme structures (Figure 2C). We then calculated the receiver operator characteristics (ROC), expressed as area under the ROC curve (AUC). The most conserved enzyme structures did belong to purine biosynthesis, several of the amino acid biosynthetic metabolic pathways (histidine, lysine, glutamate, and branched chain amino acids), as well as to some of the most central metabolic pathways, glycolysis, gluconeogenesis, and the TCA cycle (adjusted p-value < 0.05, Fisher’s exact test, Figures 2C, S2.1C). The same pathways also showed early enrichment in the ROC curve and high AUC-values (Figures 2D, S2.1D). Consistently, the same orthogroups were also enriched for the GO-slim terms associated to these pathways such as “generation of precursor metabolites and energy” and “nucleobase-containing small molecule metabolic process”. The picture that emerged for the most structurally divergent enzymes was more diverse, as they tended to be more broadly distributed across the metabolic network. However, enzymes belonging to the GO-slim term “lipid metabolic process” were strongly enriched within the orthogroups representing the 25% most structurally variable proteins. For instance, Oar1p (CR: 0.384), Tes1p (CR: 0.391), Eci1p (CR: 0.441), involved in fatty acid oxidation, and Tsc10p, (CR: 0.411) involved in the synthesis of sphingolipids, were all part of orthogroups within the top 5% quantile of structural divergence. We speculate that lipidomes of cells have a greater flexibility to adapt to the metabolic environment, as lipids contribute to the plasma membrane and secretory machinery, which interact with the outside environment. The orthogroups containing Alg13p and Alg14p were also among those displaying the most structural diversity: these two enzymes form a complex that functions as an UDP-N-acetylglucosamine transferase in N-linked glycosylation, indicating that this posttranslational modification might evolve readily.

Next, we addressed the impact of the enzymes’ molecular functions. Oxidoreductases showed the least structural diversity, while orthogroups containing hydrolases were enriched among the most structurally diverse (Figures 2C, 2E). Interestingly, the low structural divergence of the oxidoreductases could be explained by their prominent roles in central metabolic reactions, in particular, glycolysis, gluconeogenesis, and the TCA cycle. Indeed, the enrichment of oxidoreductase containing orthogroups was lost upon excluding the enriched central metabolic pathways (Figure S2.1E). Conversely, the prominence of hydrolases orthogroups among the structurally diverse cluster of lipid metabolic processes, did not fully explain their structural diversity (Figure S2.1F). We speculated that, because hydrolases do not require binding cofactors and have a low substrate specificity (Hoyos et al., 2017) they can more easily structurally diverge without being constrained by their function. One prominent set of cofactors are metal ions. Thus, we tested whether metal-containing enzymes show a lower degree of structural diversity than non-metal binders (Aulakh et al., 2024). We observed a significant difference; enzymes which did not require a metal ion showed a higher degree of structural diversity than enzymes requiring a metal cofactor (Wilcoxon-Mann-Whitney test, p < 1e-4, Figure 2F).

Other metabolic constraints on enzyme function can also act on enzyme evolution. For example, we observed that orthogroups containing enzymes with a lower number of intracellular inhibitors, a typical constraint of enzyme function (Alam et al., 2017), exhibited higher structural diversity, while orthogroups containing enzymes with a high number of inhibitors were more structurally conserved (Kendall τ = 0.22, adj. p-value < 1e-4, Figure S2.1G). For instance, Gnd2p, a central enzyme of the pentose phosphate pathway that catalyses the oxidation of 6-phosphogluconate to ribulose-5-phosphate, is inhibited by at least 45 cellular metabolites (Alam et al., 2017), and its orthogroup was one of the structurally least divergent. We thus concluded that characteristics of enzymes, such as pathway membership, dependency on cofactors and metal ions, and the number of metabolic inhibitors, constrain their freedom to structurally diverge during evolution.

### Roles of enzyme abundance and metabolic flux in structural evolution

Protein abundance has been identified as an important factor in sequence conservation (Bédard et al., 2022; Mata and Bahler, 2003). Given that enzyme expression is contingent upon an enzyme’s specific activity in relation to the cell’s metabolic demands, we hypothesised that enzyme abundance is a mechanism through which metabolism influences protein structural evolution. One of the advantages of working with *Saccharomyctina* is that many species are experimentally accessible. We obtained nine species from the UK’s National Collection of Yeast Cultures (Figure 3A) (Varma et al., 2022; Wu et al., 2017), and used proteomics to estimate their protein abundance. For this, we cultivated the cells in minimal media and harvested them exponentially in mid log-phase. We then extracted proteins, prepared tryptic digests for mass spectrometry, and analysed the proteomes using microflow chromatography, and a TimsTOF Pro mass spectrometer (Bruker Daltonics), using data independent acquisition, followed by data processing with DIA-NN (Demichev et al., 2020; Szyrwiel et al., 2024) (Figures 3A, S3.1A,B). We obtained high quality proteome data with precise concentration measurements. On average, we quantified 1,921 proteins per species, and obtained a low CV for technical replicates (median CV = 12.4%), indicating the low technical variation in the proteomes. Consistent with the observation that more abundant proteins are more conserved in other studies (Bédard et al., 2022; Mata and Bahler, 2003), our dataset revealed the relationship between protein abundance and the degree of structural diversity. Most orthogroups with a high structural diversity contained low-abundance enzymes (Figure 3B, Spearman ⍴ = 0.48, adj. p-value < 1E-27). We ascertained that the relationship between structural evolution and abundance depends on the enzyme class. Abundance and CR correlated strongly for oxidoreductases and isomerases, but this relationship was weaker for other enzyme classes, and abundance did not correlate significantly with CR (adjusted p-value ≥ 0.05) for hydrolases at all (Figure 3C). Hence, the relationship between abundance and structural diversity is dependent on the metabolic function of the enzyme.

**Figure 3:**
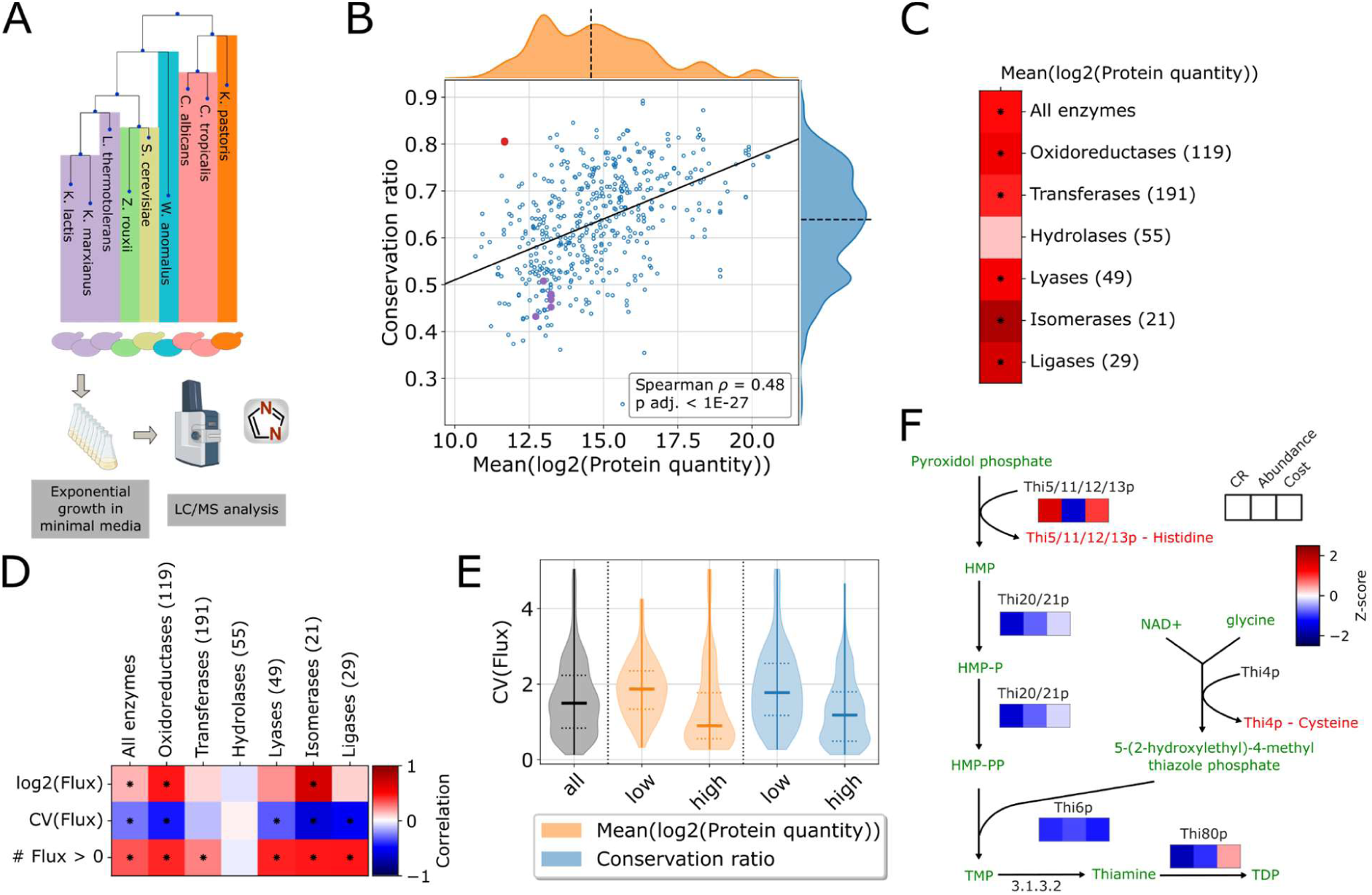
Roles of enzyme abundance and flux in structural evolution. A) Depiction of the workflow for acquiring protein abundance data. B) Mean conservation ratio versus the mean log2-transformed protein abundances from 9 of the 27 investigated species measured during exponential growth in minimal media. The orthogroup containing the Thi5/11/12/13p family is highlighted in red, other enzymes of the thiamine biosynthetic pathway in purple. The solid line indicates the best linear fit, the dashed line denotes the axis median. C) Spearman correlation between protein abundance and conservation ratio for all tested enzymes as well as broken down to the indicated enzyme class. * indicates an adjusted p-value < 0.05. The colorbar is shared with panel D. D) Correlation between different measures of predicted flux (median, coefficient of variation and number of species with flux through a given orthogroup) and conservation ratio for all tested enzymes and broken down in each column to the indicated enzyme class. For the flux median and CV, the Spearman correlation was used, and Kendall τ correlation was used for the number of species. The number in brackets indicates the number of enzymes per class. * indicates an adjusted p-value < 0.05. E) Violin plot of the CV of the fluxes for the orthogroups within the first and last quartile of conservation ratio (blue) or protein abundance (orange). F) The thiamine biosynthesis pathway is shown. Heatmaps below enzymes indicate mean conservation ratio, mean log2-transformed protein quantity, and averaged cost, with Z-scores. The Thi5/11/12/13p family and Thi4p undergo suicide reactions in which they lose a histidine and cysteine residue respectively. HMP: 4-amino-2-methyl-5-pyrimidine; HMP-P: 4-amino-2-methyl-5-pyrimidine phosphate; HMP-PP: 4-amino-2-methyl-5-pyrimidine diphosphate; TMP: thiamine phosphate; TDP: thiamine diphosphate; NAD+: nicotinamide adenine dinucleotide.

Next, we addressed the role of metabolic flux in structural diversity. We estimated metabolic flux through each pathway based on genome-scale metabolic models for each of the 26 species (Lu et al., 2021). Unexpectedly, the data revealed only a weak correlation with structural conservation (Figure 3D, Spearman ⍴ = 0.15, adj. p-value < 1E-2). Instead, we noticed a stronger interdependency between structural evolution and the variability of the flux (Spearman ⍴ = −0.27, adj. p-value < 1E-8), and that proteins with high structural diversity exhibited a fairly wide range of flux levels. In general, this paints a picture in which orthogroups with low flux variability contain highly abundant enzymes that show little structural diversity, while orthogroups with high flux variability contain less abundant enzymes that are more structurally diverse (Figures 3E, Figure S3.1C). There were exceptions to this trend, such as in the case of the orthogroup containing the sphingosine kinase Ysr3p, which was associated with a highly variable flux and showed high structural diversity, despite its relatively high abundance. In addition, we detected a dependency on the enzyme class. While structural divergence was strongly linked to the flux carried by oxidoreductases, for which all tested measures (median flux, variability of the flux, and species in which flux was present for the orthogroup) correlated with structural conservation, other enzyme classes lacked these relationships. Within orthogroups containing hydrolases especially, there was no relationship between any of the tested measures and structural diversity (Figure 3D, adj. p-value > 0.7 for all three measures).

One parameter that could influence the flux of an enzyme is its processivity (i.e. its *kcat* value). We thus estimated *kcat* values for each protein sequence in a species-specific manner (Li et al., 2022). We found only a weak relationship between structural conservation, or flux and *kcat*. However, we report that the log-transformed variability (standard deviation) of *kcat,* is higher in orthogroups that are structurally diverse (Figure S3.1D, Spearman ⍴ = −0.27, adj. p-value < 1E-8). Thus, both flux and *kcat* seem associated with structural evolution primarily through their variability, rather than their absolute values.

In parallel, we noticed that enzymes can escape the typical relationship between abundance and structural conservation due to their specific functional properties. In particular, our attention was drawn to the thiamine (vitamin B1) biosynthetic pathway. We observed that the orthogroup containing the 4-amino-2-methyl-5-pyrimidine phosphate synthase genes Thi5p, Thi11p, Thi2p and Thi13p, which occupy the same orthogroup and catalyse the same reaction, were exceptions to the overall trend and were structurally highly conserved, despite their low abundances (Figure 3B, highlighted). Notably, thiamine is extremely energetically costly to synthesize, as two of its reaction steps, including the one catalysed by these four enzymes, are catalysed by suicide enzymes that lose an essential amino acid upon catalysis (Coquille et al., 2012; Lai et al., 2012; Perli et al., 2020) (Figure 3F). In this orthogroup, evolutionary diversification is thus less constrained by abundance, but more likely by cost. We thus wondered whether metabolic cost, of which abundance is one contributing factor, might be the driving factor behind the overall relationship between abundance and structural evolution.

### Cost optimization follows a structural hierarchy

We calculated the average cost per amino acid for each protein averaged over the orthogroup using different cost metrics (Akashi and Gojobori, 2002; Barton et al., 2010; Chen and Nielsen, 2022; Craig and Weber, 1998; Seligmann, 2003; Wagner, 2005). Low-abundance, structurally diverse enzymes had more costly amino acid compositions than high-abundance enzymes (Figure 4A). Consistent with previous observations (Akashi and Gojobori, 2002; Barton et al., 2010; Raiford et al., 2008), the results indicate that high-abundance proteins evolve a less costly amino acid composition. The overall protein chain length, which is a proxy for the total cost of a protein, showed, however, only a weak correlation with the average cost per amino acid, indicating that cost optimization might be due to selection acting on specific structural elements rather than the overall protein cost, which is constrained by enzyme functional properties.

**Figure 4:**
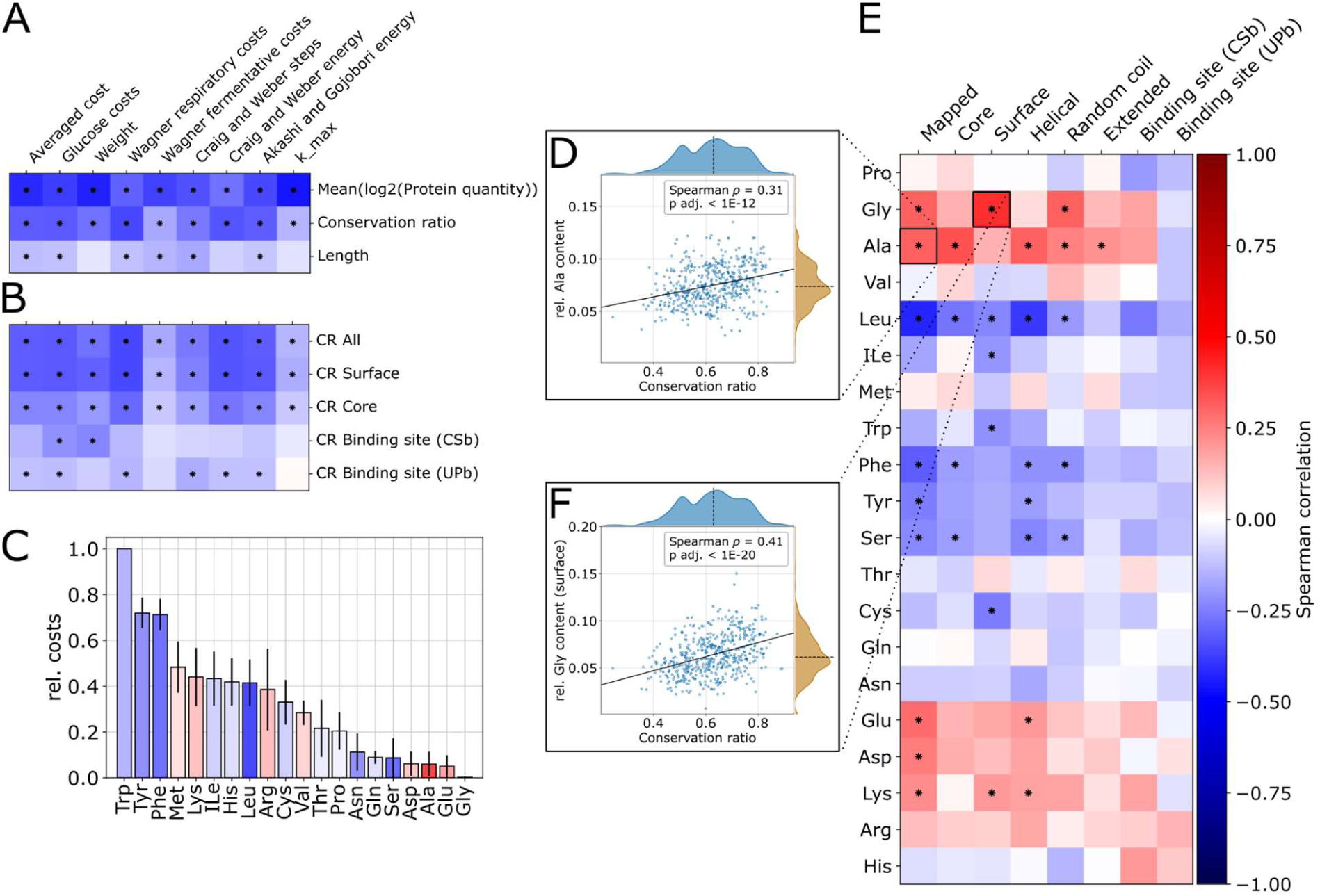
Evolutionary cost optimisation acts differently dependent on the structural element and amino acid properties. A+B) Spearman correlation of the average cost per amino acid for the entire protein versus A) mean conservation ratio, log2-transformed protein abundance and length of the protein chain and B) mean conservation ratio (CR) of selected structural features. A * indicates an adj. p-value < 0.05. C) Bar plot of the median normalised cost of each amino acid sorted in descending order. Error bars denote the MAD. Bar colouring indicates the Spearman correlation between the mean conservation ratio and the relative amino acid content of the entire protein. D) Relative alanine content of the mapped region versus mean conservation ratio. Solid line indicates the best linear fit and the dashed line denotes the axis median. E) Spearman correlation between the relative amino acid content for various structural features of proteins versus the mean overall conservation ratio. * indicates an adjusted p-value < 1E-4. F) Relative surface glycine content versus mean overall conservation ratio. Lines as in D). Plots A, B, C and E share the same colorbar shown in E.

Exploiting the protein structures at the molecular level, we were able to refine this relationship, and report that cost optimisation acts differently on structural elements, such as small-molecule binding sites, enzyme surfaces and inner structures (core residues). To identify small-molecule-binding sites, we used two strategies. First, we transferred the binding site assignments in UniProtDB to the orthogroups to account for directly coordinating residues (UPb). Second, we extracted residues in close proximity to small molecules from experimentally determined crystallographic structures from RCSB PDB, including residues within a probe radius of 5 Å, to also account for the physicochemical environment around the binding site (CSb). To distinguish surface and core residues, we used the relative amino acid-SASA and designated each residue as either a surface-exposed or core residue (Shrake and Rupley, 1973). When comparing the different structural features, we found that their amino acid composition was optimised differently in these structural elements. In general, the average cost of an enzyme’s core residues was higher than the cost of its surface residues, whereas the cost of its binding site residues was more variable. Among surface residues, we also observed increased costs for membrane-bound proteins and short proteins (<100 amino acids), such as Qcr8p or Kti11p in *S. cerevisiae*, which might be part of larger complexes (Figure S4.1A).

Studying cost optimization in more detail, we observed a hierarchy of structural evolution. Enzymes with more conserved surfaces tended to be less costly than enzymes with more structurally variable surfaces. This effect, although less strong, was also present for core residues, and it was drastically diminished for binding sites, indicating that the pressure to constrain costs is exerted on the surface and core, but not for enzyme binding sites, to prioritise biochemical function (Figure 4B). Moreover, the more expensive aromatic amino acids, such as phenylalanine and tyrosine, were more frequent in the structurally variable and low-abundance enzymes, than structurally conserved, high-abundance enzymes. The latter contained higher amounts of the least expensive amino acids, glycine, glutamate and alanine (Figure 4C, D and S4.1B, C). Leucine, lysine and serine, however, showed exceptions to these trends. While leucine and lysine add intermediary costs, one was more abundant and one was less abundant amongst the structurally divergent orthogroups, while low-cost serine was found more often among the structurally divergent structures.

We broke this analysis down to the molecular level (Figure 4E and S4.1D) and determined that evolutionary constraints act differently on the different enzyme substructures. For 11 of the 20 canonical amino acids, a significant association (adj. p-value < 1E-4) was detected in at least one structural feature. For instance, surfaces and coil regions of highly structurally conserved enzymes had higher relative glycine contents (Figure 4F), while the core and helical regions had higher relative alanine contents. *Vice versa,* the glycine content in the core and alanine content on the surface were not significantly different between conserved and variable structures presumably, because these small and inexpensive amino acids can replace many other amino acids, and their sterical and physicochemical properties are more suited to either the core or surface, respectively. Furthermore, the surfaces of the variable enzyme structures had higher cysteine contents, a correlation not found in other structural elements. This is presumably because cysteine often performs specific dedicated functions related to its high reactivity, such as catalytic activity or stabilisation via disulfide bridges. A recent study of human proteins observed that cysteine surface accessibility was related to the dedicated function of cysteine (White et al., 2023). There were no significant associations between the presence of any amino acid in binding sites and the CRs of the enzymes. Taken together, these results show that, away from binding sites, structural features influence the way in which highly conserved enzymes optimise costs. Within binding sites, costs are not optimised, which supports the notion that structural evolution occurs while preserving the biochemical reaction mechanism of the enzymes.

### Hierarchy of evolution of structural features

Notably, across all orthogroups, we detected less structural diversity in core enzyme regions than on surfaces (Figure 5A). At the level of the secondary structure, most helical portions (e.g. ɑ-helices) varied more than β-sheets. To our surprise, they were also more variable than mapped random coils, which are assumed to have a higher dynamic flexibility (Figure 5B, Figure S5.1A, Supplementary Note 3). As indicated by cost analysis, the binding sites were not only the most conserved regions (Figure 5C, Figure S5.1B), but orthogroups that contained less variable enzyme structures were also significantly enriched for possessing a known binding site (AUC = 0.62/0.66, Figure 5D). Surprisingly, for Fas2p (N = 1887 amino acids in UniProtDB) with both binding site definitions (N = 21/95 amino acids), the binding site seemed to be more variable than the overall protein. This is not the case for most other orthogroups, in which the enzyme as a whole evolved faster than the binding site (Figure S5.1 C, D), reflecting the conservation of enzymes’ catalytic functions.

**Figure 5:**
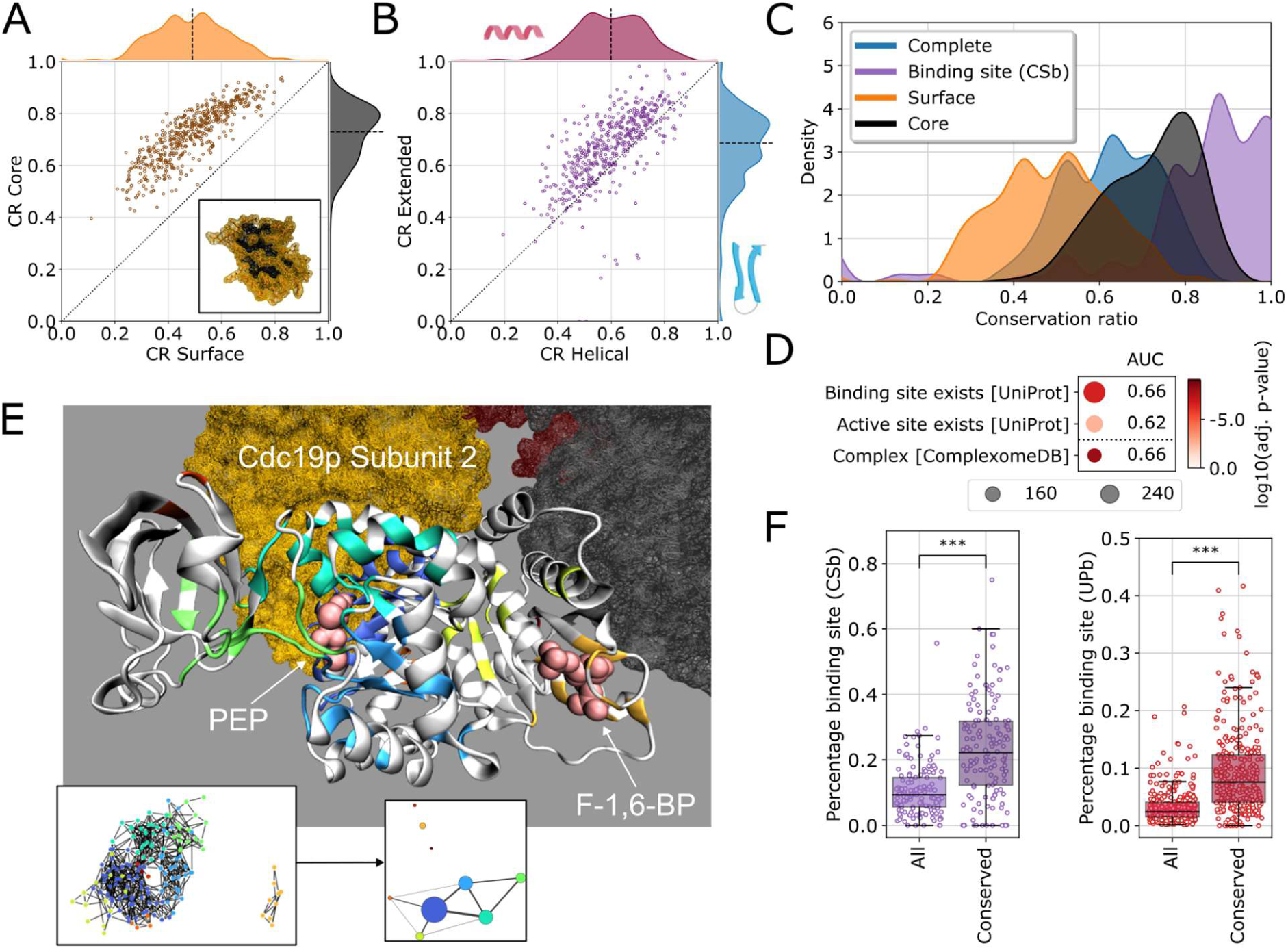
Hierarchy of evolution of structural features. A) Mean conservation ratio of the core versus mean conservation ratio of the surface. Dotted line denotes the identity line and the dashed line denotes the axis median. In the lower right a depiction of core (black) and surface parts of the protein (orange) based on relative solvent accessibility surface area (SASA) on the example of 5-Formyltetrahydrofolate cyclo-ligase is shown. B) Mean conservation ratio of the extended versus mean conservation ratio of the helical parts. Lines as in A). C) Distribution of the mean conservation ratio for all mapped residues (blue), for just the core (black) or surface residues (orange) and for the binding site residues (purple). For the binding site, the crystal-structure-based definition (CSb) is shown. D) Enrichment of orthogroups containing the 25% most conserved enzymes for structural features and catalytic properties. Dot size indicates the number of metabolic enzymes associated with the relevant list in our selection, colour indicates the adjusted p-value. E) Depiction of the conserved regions in Cdc19p. Each non-white region denotes a fully conserved cluster of amino acids. The spheres indicate the bound ligand (PEP, associated clusters: dark blue, light blue, green, and cyan) and activator (F-1,6-BP, associated cluster: orange) (crystal structure: 1A3W (Jurica et al., 1998)). Other subunits in the homotetramer are highlighted in yellow, red and black. In the lower left inset, the clustered network is shown coloured according to the cluster label. Each node is a fully conserved residue, and an edge is assigned between two residues when their Cɑ-atoms are within 10 Å of one another. In the lower right inset, a summary of the clustered network is shown. The node size indicates the cluster size, the edges indicate cluster connectivity and the edge width indicates the number of connecting edges from the original network between the two clusters. F) Percentage of either all or only fully conserved residues that are identified as binding sites for both the CSb definition (purple) or UPb definition (red) of binding sites. The boxes denote the first and third quartile, the whiskers extend up to 1.5-times the interquartile range. *** indicates a p-value < 1e-4, Wilcoxon signed-rank test.

To obtain an orthogonal view we identified spatially proximate, highly-conserved regions within enzymes, which were likely to have been conserved to perform important functions for the cell. To extract those regions, we clustered a network of fully conserved residues, resulting in a median number of 4.00 (IQR: [2.00, 6.00]) clusters with a median size of 18.55 amino acids (IQR: [14.00, 23.64]) per cluster (see Table S5.1 for a more detailed analysis). An example is pyruvate kinase (Cdc19p of *S. cerevisiae*). We located the activation site binding the allosteric activator fructose-1,6-bisphosphate (F-1,6-BP) and the binding site for its substrate phosphoenolpyruvate (PEP), however, we also observed a highly conserved, symmetric interaction surface facing another protein subunit (dark blue in Figure 5E).

Comparing all orthogroups, we detect over 2-fold more substrate/ligand binding site residues within these fully conserved clusters (Figure 5F). However, 27% (CSb) or 50% of these clusters (UPb) did not overlap with known binding sites and thus the remaining clusters might serve some other purpose, e.g. provide overall stability or interaction sites for other biomolecules, unknown ligands, or other protein subunits as shown in the dark blue cluster in Cdc19p. Consistent with this observation, enzymes with more physical protein-protein interactions (Figure S5.1E; Kendall τ = 0.32; adj. p-value < 1E-26) and enzymes involved in protein complexes (ComplexomeDB, Figure 5D) (Benschop et al., 2010) were more highly conserved.

### Structural divergence due to positive selection in the electron transport chain

Finally, we addressed whether an increase in structural diversity is explained by evolutionary selection. We focused on the orthogroups that contain the most variable enzyme structures and asked to which degree their increased structural diversity represents drift or positive selection by comparing the structural diversity detected within the orthogroups with dN/dS values, that in evolutionary genomics are used to distinguish selective neutrality (dN/dS of ∼1) from positive (> 1) and purifying selection (< 1) (Kimura, 1977; Miyata and Yasunaga, 1980). We calculated dN/dS ratios based on all-vs-all alignments to initially calculate a single rate per orthogroup (M0 model) to assess the extent of this (Álvarez-Carretero et al., 2023; Jeffares et al., 2015; Zhang and Skolnick, 2005). Although we obtain an average dN/dS of < 1, indicating overall purifying selection occurred for each orthogroup, we report substantial differences depending on the enzymes. Relatively high values of dN/dS may indicate orthogroups with higher proportions of residues under neutral drift (dN/dS = 1) or positive selection (dN/dS > 1). As our selected orthogroups spanned a large evolutionary timespan (400 my), the calculated dS values for some of the orthogroups were saturated, which could have skewed the dN/dS calculations. At the expense of removing some low conservation orthogroups, we removed orthogroups with an average dS value of > 3 (Figure S6.1A, B). These were enriched for several GO terms related to glycosylation. While orthogroups with dS < 3 were enriched for terms related to central carbon metabolism (Figure S6.1C). For the remaining orthogroups, we performed enrichment analysis for the first and last quartiles of the dN/dS values (Figure 6A). For orthogroups in the first quartile of dN/dS values, glucose fermentation and ETC pathways were enriched, as well as the GO slim terms “oxidoreductase activity”; “monocarboxylic acid metabolic process”; the molecular function term “transmembrane transporter activity”; and the cellular compartment terms “membrane”, “mitochondrion”, and “mitochondrial envelope”. No terms or pathways were significantly enriched in the orthogroups of the last quartile of dN/dS values. Although no enzyme class was significantly enriched, there were differences at the level of individual enzymes with specific oxidoreductases and isomerases, as well as specific enzymes of the the GO term “lipid metabolic process”, having high dN/dS values (Figure S6.1D, E).

**Figure 6:**
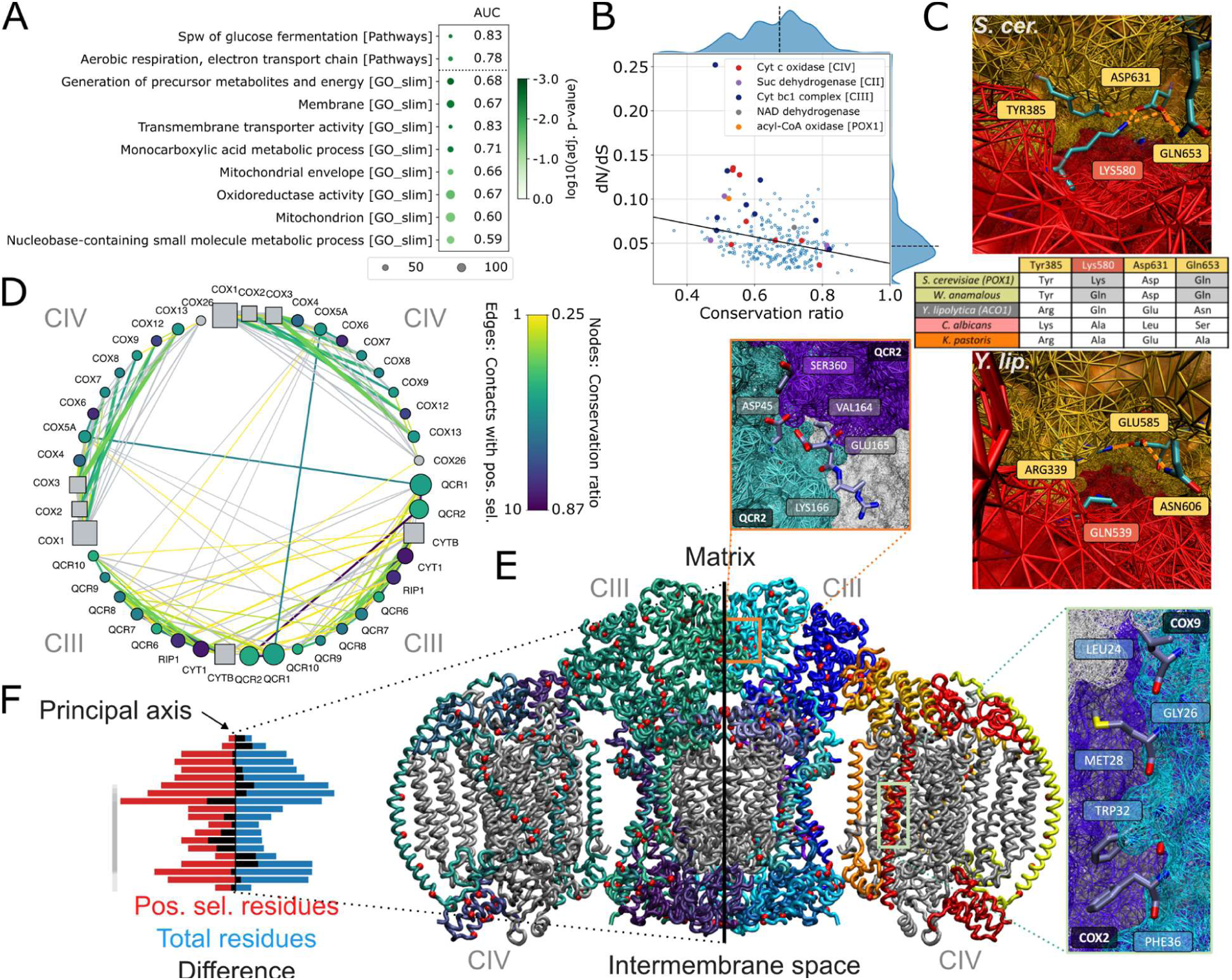
Pox1p and several proteins of the Electron Transport Chain are among proteins with high structural divergence and higher levels of positive selection. A) Enrichment of the orthogroups containing enzymes with the 25% highest dN/dS values with respect to metabolic pathways and GO slim terms. For the lowest 25% dN/dS values, no significant enrichments could be detected. Dot size indicates the number of enzymes in *S. cerevisiae* associated with the pathway or GO term from our dataset, colour indicates the adjusted p-value. In addition, the AUC is reported. B) dN/dS value versus mean conservation ratio per orthogroup. Orthogroups that contain proteins of the electron transport chain are highlighted, as well as the orthogroup containing *S. cerevisiae* Pox1p. The solid line indicates the best linear fit and the dashed line denotes the axis median. C) Structure of the fatty-acyl coA oxidase homodimer interface obtained by using the crystal structure of the *Y. lipolytica* AOx protein, ylACO1 (bottom), (PDB:5Y9D (Kim and Kim, 2018)). This crystal structure was also used as a reference to superimpose *S. cerevisiae* Pox1p structure from Alphafold2 (top). The positively selected amino acids Lys580 and Gln653 as well as the coordinating amino acids Tyr385 and Asn631 are highlighted. The red and the orange surfaces denote different protein chains of the homodimeric complex. The inset table denotes the residues present for various orthologs. Grey shading in the table denotes residues and branches that were under positive selection. D) Connectivity network of the cytochrome-c-oxidase/cytochrome bc1 complex (PDB: 6HU9 (Hartley et al., 2019)). The node size indicates the length of the protein chain. Square nodes denote mitochondrially encoded genes. The nodes are coloured with respect to the conservation ratio, where grey denotes missing orthogroups in our data selection. The edges indicate physical interactions between the proteins and edge width indicates the number of contacts between the surfaces of the two protein chains. The edges are coloured according to the number of positively selected amino acids from either protein chain on the surface between the two protein chains. E) Crystal structure of the cytochrome-c-oxidase/cytochrome bc1 complex. The structure consists of a homodimer made up of two ubiquinol cytochrome c reductase (complex III) complexes surrounded by two cytochrome c oxidase (Complex IV) complexes (Smith et al., 2012). Protein chains are coloured with respect to their conservation rate (left part) or chain ID (right part). Red dots denote positions where positive selection was detected in at least one branch. The complex is symmetrical across the black line. In addition, two interfaces are shown in detail, highlighting positively selected residues between the two surfaces. F) Histogram (20 bins) of the normalised total number of amino acids (blue) and normalised number of positively selected amino acids (red) along the black line of symmetry in E) per bin. The black bins indicate the difference between the red and the blue bins. The grey line on the left of the diagram indicates the presence of mitochondrially encoded protein chains.

Our interest was triggered by the orthogroups that were highly structurally diverse (CR ≤ 0.6) yet had higher dN/dS values (> 0.06), indicating a higher proportion of residues under positive selection. This group included several proteins of the ETC (Figure 6B and Figure S6.1F), such as complex III (in 6 of the 9 orthogroups) and complex IV (in 4 of the 8 orthogroups) as well as the fatty acyl coA oxidase Pox1p that is part of the fatty acid β-oxidation pathway. To investigate positive selection in these orthogroups in more detail, we obtained dN/dS estimates at the amino-acid level for the orthogroup containing Pox1p as well as for orthogroups of the ETC (Yang, 1998; Yang et al., 2005; Yang and Nielsen, 1998; Zhang et al., 2005).

### Positive selection in structurally diverse orthogroups at protein-protein interaction surfaces

In the Pox1p orthogroup’s protein tree, we identified 15 branches, some of which were nested, showing evidence of positive selection (Figure S6.2A). Within the branch encompassing *S. cerevisiae* and *Wickerhamomyces anomalous*, 15 residues showed signatures of positive selection, including residues aligning with Lys580, and Gln653 in *S. cerevisiae*. After projecting these residues onto an experimentally determined protein structure of the homodimer in *Y. lipolytica* (Kim and Kim, 2018), we report that, under physiological pH, positively charged Lys580 appeared to physically interact with the negatively charged Asp631 across the homodimer interface in this branch, while Gln653 and Tyr385 stabilise Asp631 (Figure 6C), forming a stabilising ionic and hydrogen-bond interaction. These residues differ from their counterparts outside this branch (Figure S6.2B, Table S6.1). For instance, in the branch containing *Y. lipolytica* ylAOX1, these interactions across the homodimer interface were absent (Figure 6C) and were replaced with stabilising interactions between Glu585 and Arg339 and Asn606. We speculated that these residues may have been selected in the common ancestor of *S. cerevisiae* and *W. anomalus* for stronger interactions at the homodimer interface, as the interaction occurs twice on the surface, which could enhance fitness even more.

For the ETC, we obtained up to 11 branches with evidence of positive selection with an average of 2.3 and up to 30 positively selected residues per branch for each orthogroup (Figure S6.3 A, B). To determine if these positively selected residues also appear on protein-protein interaction interfaces, we focused on the ETC ubiquinol cytochrome c reductase/cytochrome c oxidase supercomplex (CIII_2_-CIV_2_ mitochondrial respiratory supercomplex, PDB: 6HU9 (Hartley et al., 2019). The supercomplex spans the membrane which separates the mitochondrial matrix from the intermembrane space. For each protein chain, we investigated structural conservation, protein length, connectivity, number of interactions, and number of positively selected residues between the surfaces (Figure 6D). We report several instances, where a structurally diverse orthogroup showed positively selected residues at interaction surfaces. Of the 157 unique positively selected residues identified, 96 were on protein-protein interaction surfaces. Within the complex, most positively selected residues were detected on the homo-dimeric Qcr2p-Qcr2p and the Qcr2p-Qcr1p interfaces, and between complexes, most were at the connection interface Cox5Ap-Qcr1p (Figure 6E, Figure S6.3C, Inset). Of note, the dimeric Qcr2p-Qcr2p interface contained five positively selected residues in both the branch containing orthologs from *S. cerevisiae* to *W. anomalus* and the branch containing orthologs from from *S. cerevisiae* to *A. rubescens* that form a cluster on the interface which amplifies any effects due to the dimeric characteristic of the interface (Figure 6E, Figure S6.3D).

The orthogroups containing mitochondrial encoded ETC complex did not pass our inclusion criteria, meaning that these orthogroups were underrepresented in the interface analysis. Nonetheless, there were four positively selected points of contact between the nuclear encoded Cox7p and the mitochondrially encoded Cox3p, and five positively selected points of contact between the nuclear encoded Cox9p and the mitochondrially encoded Cox2p (Figure 6D, 6E). We hypothesised that positive selection is stronger for these nuclear encoded proteins, as the mitochondrial genome has been shown to have a higher mutation rate than the nuclear genome in more recently diverged branches of budding yeast (Christinaki et al., 2022), which could also explain their high structural diversity. To assess this, we compared the total number of amino acids present along an axis of symmetry perpendicular to the membrane to the number of amino acids along that axis that showed positive selection. In the central part of this axis, where there was a higher proportion of mitochondrially encoded proteins, there was a greater number of positively selected residues, although the difference in density distributions of all residues versus that of positively selected residues was not significant (Figure 6F, two-sided, two-sample Kolmogorov-Smirnov test, p < 1e-1).

## Discussion

Building on decades of painstaking work to experimentally determine protein structures, deep-learning based computational tools have reached a level of maturity at which they can predict high quality protein structures from sequence data (Ahdritz et al., 2023; Baek et al., 2021; Jumper et al., 2021; Lin et al., 2023). Although structural predictions still have limitations (Terwilliger et al., 2024), tools such as Alphafold2 have, for the first time, made structures available systematically at large scale and across species barriers. Having structures available at this scale generates new possibilities to address higher order biological problems, such as the mapping of evolutionary constraints acting on proteins over time. Recently, predicted structural information was used to classify and identify structural folds across the range of biological variation (Barrio-Hernandez et al., 2023; Durairaj et al., 2023; Pavlopoulos et al., 2023). For other problems, we are currently only beginning to exploit these newly added possibilities for making biological discoveries.

In this study, we combined comparative evolutionary genomics with structural biology to map the metabolic constraints that have affected protein evolution. In order to achieve a sufficient statistical resolution and to have sufficient data at hand in order to link all genomic, structural, proteomic and metabolic evolution, we focussed on the *Saccharomycotina* subphylum. The information used in our study included not only well-annotated genome sequences, on which basis we predicted enzyme structures using Alphafold2 (Jumper et al., 2021) but also enzyme functional annotations, genome scale metabolic models, and proteomic data, accrued by the research community, due to the longstanding importance of this subphylum in biotechnology, microbial ecology, infection biology, and as a model organism. The 11,269 enzyme structures analysed in our study stem from 27 selected genetically and metabolically divergent species, covering most metabolic pathways, GO terms, and enzyme classes present in this exquisitely well studied subphylum.

We asked if metabolic properties that differ between species, pathways, and enzyme classes can explain the range of structural diversity observed in the different orthogroups. In this way, we identified the metabolic constraints that influence structural evolution at different biological scales, from the level of the organism down to the pathway, to individual enzymes, and to enzyme substructures. We also identified pathways, enzyme classes, and, in some instances, structural elements that contribute to these relationships.

At the scale of the organism, we identified enzymes that were either more structurally conserved or more variable in species that can ferment glucose versus species that have an oxidative metabolism, the two major metabolic niche specialisations in eukaryotic cells. Several of these enzymes were shown to be functionally related to glucose fermentation, though some were not, as multiple major phenotypic changes can occur in the same clade. Nonetheless, this indicates that enzyme structural evolution is often a consequence of broader rearrangements, i.e. enzyme structural diversity is dependent on metabolic niche adaptations.

A more detailed picture of this process was obtained at the metabolic pathway scale. At this scale, we revealed that orthogroups of enzymes involved in central carbon metabolism, as well as oxidoreductases and metal binding enzymes, were the least structurally divergent, while those of lipid metabolism, a more peripheral metabolic pathway, and enzyme classes, such as hydrolases, diversified structurally. Overall, our data suggest that the evolution of an enzyme depends on the evolutionary constraints of the pathway it participates in, the type of reaction it catalyses, and other cellular microenvironmental factors such as the number of cellular metabolites that inhibit the enzyme or the dependency of an enzyme’s reaction mechanism on cofactors, such as metal ions. We thus concluded that function, reaction mechanism, and the metabolic microenvironment of an enzyme constrain its structural evolution.

At the enzyme level, we found that, while abundance constrains structural divergence, it can be trumped by overall metabolic cost. For example, we noted that enzymes of the thiamine metabolic pathway (Thi5/11/12/13p), which carries major metabolic costs due to the involvement of a suicide enzyme that is irreversibly inhibited upon catalysing a single reaction, remain highly structurally conserved despite their low abundance. We also compared metabolic flux levels with structural divergence and found that, while there is some relationship between flux and structural variability, a stronger relationship is obtained considering the variability of the flux, and that the effect of flux variability can override the effect of abundance on structural divergence. For instance, the enzyme Ysr3p showed high structural diversity despite its relatively high abundance, but it also showed high variability in flux. Remarkably, these relationships also depend on the metabolic function of an enzyme. Hydrolases, the least conserved enzyme class, are also relatively independent of the metabolic microenvironment, as they are cofactor-independent, and have largely escaped the selective pressure of these relationships.

Overall, at the enzyme substructure scale, our analysis revealed a hierarchy of structural diversity, with surface structures being most diverse and small-molecule-binding being the most structurally conserved. Enzymes with more conserved surfaces tended to be less costly than enzymes with more structurally variable surfaces, while the most conserved spatially co-located regions of proteins were enriched for small-molecule-binding sites. Essentially, this hierarchy of substructural conservation is consistent with a model in which metabolic enzymes evolve alongside the chemical topology of the metabolic network, with structural components involved in catalysis changing the least. Notably, we report clusters of high structural conservation that lack any functional annotation. It is worth speculating that many of these might represent thus far unmapped protein-protein interaction sites. Comparative structural biology could thus be a tool to identify sites of enzymes of unknown function.

Also at the enzyme substructure scale, we gained insight into the way evolution achieves cost optimization. Binding sites did not appear to optimise cost, presumably because they must optimise enzyme catalytic function regardless of the expense of their constituent amino acids. Furthermore, cost is optimised in structural features differently. For instance, alanine is specifically enriched in the cores of highly conserved enzymes, while glycine residues are enriched on their surfaces. We suggest that future cost metrics should factor in substructural information.

Finally, we assessed whether the higher structural divergence of some enzymes represents a lack of constraints (allowing genetic drift) and greater freedom to evolve, or instead whether it represents adaptive evolution. In some enzymes we observed regions that exhibit evidence of positive selection. After studying representative examples, such as Pox1p and respiratory chain complexes, in more detail, we concluded that some regions showing positive selection represent protein-protein interaction sites which have co-evolved in enzyme complexes. Thus, identifying highly conserved regions as well as signals of positive selection in 3D structures could be developed into new strategies for enzyme functional annotation, and to understand and engineer molecular networks. For example, combining the presented framework with (I) binding site prediction data (Tubiana et al., 2022), protein small-molecule interaction prediction data (Abramson et al., 2024; Ferreira et al., 2015; Hekkelman et al., 2023) and domain structural similarity information (Barrio-Hernandez et al., 2023) could help locate binding sites; with (II) protein-protein interaction studies (Michaelis et al., 2023) and protein complex prediction algorithms (Abramson et al., 2024; Akdel et al., 2022; Bryant et al., 2022; Evans et al., 2022; Mirdita et al., 2022) could help identify protein-protein interaction surfaces; and with (III) metabolic engineering techniques (Nielsen, 2001; Sharma et al., 2021) integrated with the identification of functional intrinsically disordered regions utilising molecular dynamics (Tesei et al., 2024), it could be used to obtain information for protein editing. These approaches could help realise the full potential of systematically applying structural information to help link enzyme structure to metabolic function.

In summary, in this study, we combined comparative evolutionary genomics with structural biology by leveraging Alphafold2’s ability to predict enzyme structures systematically and across species barriers. We mapped biochemical constraints on protein evolution in the *Saccharomycotina* subphylum, by studying structural divergence within 11,269 enzyme structures from 27 species. We found that metabolic properties influence structural diversity across species, pathways, and enzyme classes. Enzyme structural evolution correlates with metabolic niche adaptations, with central carbon metabolism enzymes being more conserved and lipid metabolism enzymes showing greater diversity. We also observed that enzyme abundance and metabolic cost constrains structural divergence, while flux variability also plays a significant role. At the substructural level, small-molecule-binding sites are the most conserved. These findings illuminate the relationship between enzyme function, metabolic environment, and structural evolution, offering new strategies for enzyme annotation and metabolic network engineering.

## Supplementary Notes

### Note 1: Choice of *Saccharomycotina* yeasts for addressing metabolic evolution

The yeast subphylum *Saccharomycotina* shared a common ancestor around 400 million years ago (MYA). Unlike bacteria, however, there is much less horizontal gene transfer which would complicate comparisons across different orthogroups of proteins. At the same time, they are still characterised by a remarkable metabolic diversity, including gene losses preventing growth on specific substrates (Opulente et al., 2024; Shen et al., 2018), a major whole-genome hybridization that corresponds with the emergence of the ability to ferment in the presence of oxygen (Crabtree effect) (Hagman et al., 2014; Hagman and Piškur, 2015; Marcet-Houben and Gabaldón, 2015), and limited horizontal gene transfers from other yeast and bacteria that confer new metabolic capabilities (Gonçalves et al., 2018; Gonçalves and Gonçalves, 2022; Kominek et al., 2019; Marsit et al., 2015). The clade has been comprehensively sequenced and characterised at the molecular and metabolic level (Kurtzman et al., 2011; Opulente et al., 2024; Riley et al., 2016; Shen et al., 2018; Steenwyk et al., 2022; Wolters et al., 2023; Wu et al., 2017), and includes the most prevalent human fungal pathogen *Candida albicans,* and several industrially important yeast species (*Kluyveromyces marxianus, Komatagella pastoris, Yarrowia lipolytica)* as well as the model single celled eukaryote and food and beverage industry workhorse, *Saccharomyces cerevisiae.* Previous work has integrated phenotypic and genomic evidence to understand the evolution of metabolism in the yeast subphylum (Opulente et al., 2024; Shen et al., 2018), including reconstruction of individual genome scale metabolic models for hundreds of species (Li et al., 2022; Lu et al., 2021).

### Note 2: Limitations of AlphaFold2 in relation to our study

It has been shown that the quality of AlphaFold2 predictions can vary depending on the predicted protein. Although in many cases there is a good agreement with experimental data, in some cases inaccuracies on the global or local level have been observed. Examples are domain displacements or inaccurate side chain orientations (Terwilliger et al., 2024). We observe a few rare instances of domain displacement in our data, however, since we observe a median mapping ratio of 87.4% and we detected this behaviour only sparsely, it should not strongly influence our analysis. Since our mapping and analysis protocols are based mainly on the backbone configuration, inaccuracies in the side chains should also not influence the presented results.

In this manuscript we assigned the binding site using evidence from crystal structures in the PDB. This was done because in Alphafold2 structures do not explicitly include substrates, cofactors, other protein chains and other coordinating subunits. However, since AlphaFold2 is a deep-learning framework that utilises existing structures which contain these additional molecules, it implicitly incorporates information about their presence, but the nature and extent to which that information is included is lost (Jumper et al., 2021). The very recent publication of Alphafold3 allows the inclusion of such interactions and could be incorporated in future work (Abramson et al., 2024). Additionally, Alphafold2 does not accurately capture structural information from (intrinsically) disordered domains (Tunyasuvunakool et al., 2021) because the dynamical properties of the protein are not included in its training. Dynamic disordered domains, although present, (Chakravarty and Porter, 2022; David et al., 2022; Ruff and Pappu, 2021) are less common in metabolic enzymes than they are in some other classes of proteins (Bondos et al., 2021; DeForte and Uversky, 2017), so by focusing on metabolic proteins, we avoid this issue to some extent.

### Note 3: Observations that require known structures

In this work, we aimed to investigate structural changes that might affect evolution on a large scale which might not be visible when analysing protein sequences, or the smaller set of known experimental structures alone. We saw that surface residues were less conserved than interior residues. Reasons for this might be multifold: increased steric hindrance due to close packing of the protein, stabilising interactions, protein folding and solubility. In addition, surface residues might be more prone to mutational drift if they don’t fulfil a functioning, coordinating or stabilising purpose (Choi et al., 2009). Also, for extended structures and random coils a higher conservation than for helical regions was observed. This may be partially explained by a higher number of interactions between the residues of the extended structures and random coils than within helical structures. In the case of random coil structures, we also only took those into consideration that could be mapped to the respective reference structure and thus seem to have, at least in the prediction, a defined spatial arrangement.

The structural information that we had for our orthogroups also allowed us to better understand relationships between conservation and relative amino acid content, a sequence-based factor. We were able to further analyse these relationships in the context of different structural features to determine how those features contributed to the relationship. We find, for instance, that the correlation between alanine content and conservation is more specific to helical regions, while the correlation between glycine content and conservation is more specific to random coil regions. For alanine it might be explained by its helix formation properties (Chakrabartty et al., 1994) while for glycine it might reflect the increased backbone flexibility due to the lack of a side chain (Ho and Brasseur, 2005). Also, for core and surface residues differences could be observed. The fact that the correlation between CR and amino acid content is stronger for alanine within inner residues and for glycine within outer residues might be due to the more hydrophobic nature of alanine compared to glycine (Kyte and Doolittle, 1982). This observation is consistent with a notion of highly conserved proteins optimising for cost in external residues by replacing more expensive amino acids with less expensive glycines when possible and doing the same with alanines for internal residues. We also see a surprising anticorrelation between leucine content and overall conservation, which is more prevalent when considering helical and coil regions than when considering extended regions and might be a direction for future research.

Also for cysteine an anticorrelation with CR is observed only for outer residues, likely since it is low in overall abundance (Figure S4.1B) and typically has specific dedicated functions, related to its reactive nature, such as catalytic activity or stabilisation via disulfide bridges (White et al., 2023), in the protein. In the human proteome, for example, a recent study observed that cysteine surface accessibility was related to the dedicated function of the cysteine (White et al., 2023). One way of understanding this observation is to consider that cysteine on the outer residues of a protein may cause an increased burden on the cell for expressing the protein, thus driving a negative correlation with conservation as selection tends to minimise that burden. This may be due to selection against spontaneous, unintended reactions of surface-exposed cysteines in more conserved proteins (Marino and Gladyshev, 2010). For other amino acids, such as proline, no significant correlation was observed, and, for proline, a median correlation around zero was obtained. This could possibly be related to the fact that proline limits not only its own backbone flexibility, but also that of its preceding amino acid (Vitalini et al., 2015) and thus might be important for a structure’s overall spatial arrangement.

## Methods

### Orthogroup and Species Selection

We selected a range of 26 sequenced budding yeast species that span the diversity of the *Saccharomycotina* subphylum, as well as the model fission yeast *Schizosaccharomyces pombe,* as an outgroup to root the tree. Nine of the species were selected, including the model budding yeast, *S. cerevisiae*, were identified in (Wu et al., 2017) as a phylogenetically diverse set of yeast species representing the NCYC strain collection. We also chose the medically relevant commensal and pathogenic yeast, *C. albicans* among 15 other species chosen to represent all the major clades of the subphylum as described in (Shen et al., 2018) (Figure 1A, Table S1.1).

Orthogroups were assigned as per orthoMCL clusters from (Shen et al., 2018). In order to select orthogroups for the study, we began with genes assigned to yeast pathways per a yeastmine query mapping pathways to genes executed on 07 Oct 2021 (Wong et al., 2023). We used 517 of the 555 genes in the pathway database (“YeastPathways Database Website Home,” 2023). The missing 38 genes came from 51 pathways not included in our yeastmine query due to a data error that we reported to SGD and which has been fixed. From these 517 genes we obtained 445 orthogroups as some orthogroups contained paralogs of some pathway genes. From those orthogroups, we kept 426 orthogroups that had genes present in at least 40% of the species. Those 426 orthogroups contained 534 genes in *S. cerevisiae* and, of those, 499 genes (90%) were associated with 224 of the 229 (98%) pathways in the yeast pathways database.

### Structural predictions using AlphaFold2

We downloaded structures for *S. cerevisiae*, *C. albicans* and *S. pombe* from the AlphaFold Protein Structure Database (Jumper et al., 2021) on 27 Apr 2022 [*S. cerevisiae* (n = 534)*, S. pombe* (n = 375) and *C. albicans* (n = 392)]. For the other species we predicted protein structures based on sequences using Alphafold2. We attempted to obtain protein sequences from uniprot for the 9 species identified by (Wu et al., 2017) and for which we had proteomics data. As the genes name from (Shen et al., 2018) and that from uniprot did not correspond, we matched proteins by calculating pairwise protein similarity between each gene identified in (Shen et al., 2018) and every gene in the uniprot proteome using the biopython function pairwise2.align.globalms with the following parameters: match_points = 1, mismatch_points = −1, gap_open = −.5, gap_extension = −0.1, and penalize_end_gaps=True. Where there was a similarity score above 75, or where there was a similarity score above 60 with a difference of 8 or greater above the next highest similarity score, we used the uniprot protein sequence. Where these conditions were not met, we used the protein sequence from (Shen et al., 2018). For species not studied in (Wu et al., 2017) we used protein sequences from (Shen et al., 2018).

We provided 10,545 sequences as input to Alphafold2 version 2.0.1, installed on the Berzelius computing infrastructure and using the default settings with the full BFD database. The structure with the best pLDDT-score of the 5 output models was used as our final structure. We were unable to calculate structures for 577 sequences, and the distribution of these sequences was skewed in terms of sequence length, species, and orthogroup (Tables S1.2-3). For the sequences for which we couldn’t calculate structures, 20.8% had a length greater than 1800 residues, compared to only 0.6% of the sequences for which we could calculate structures. This length dependence was apparent in the bias in the unpredicted structures towards specific orthogroups. 29.3% of uncalculated structures were in 7 orthogroups for which structures could not be generated in over half of the sequences of the orthogroup. This included two orthogroups for which no structures could be generated for any of the sequences (27 sequences from OG1710 which includes the *S. cerevisiae* proteins Hfa1p and Acc1p, and 24 sequences from OG1869 which includes the *S. cerevisiae* gene Fox2p). For these 7 orthogroups, 55.6% of their sequences were longer than 1,800 residues (Table S1.2).

There were also more unpredicted structures in *Hanseniaspora osmophelia*, *Zygosaccharomyces rouxii*, and *Yarrowia lypolitica* relative to other species (163, 131, and 48 unpredicted structures respectively), comprising 59.3% of the unpredicted structures. Length did not appear to be a strong factor in preventing the prediction of these sequences, as only 5.0% of them had length greater than 1,800. In sum of the 577 structures, we could not calculate, 88.4% were either from one of these three species, in one of the 7 most challenging orthogroups, longer than 1,800 residues, or a combination of those factors. Following structure prediction, we combined our predicted structures with those predicted in the AlphafoldDB from our model organisms and carried 11,269 structures from 424 orthogroups forward for further analysis.

For the pLDDT, especially for the N-terminal region low values were obtained. One reason might be that the annotation of translational start sites can be difficult to predict from genomic sequences, and alternative canonical start codons as well as alternative non-canonical start codons can be used for various genes (Eisenberg et al., 2020). Thus it is possible that for some genes the annotated N-terminal region is not expressed. Also both N-terminal and C-terminal regions of a protein often contain localization sequences, regulatory domains, or domains that modify protein-protein interaction (Almagro Armenteros et al., 2019; Sharma and Schiller, 2019). These are often unstructured and may be cleaved and degraded, therefore these regions may not be selected for structural stability.

### Multiple structure alignment and Orthogroup refinement

For each of the orthogroups, multiple structure alignments were carried out initially on the 424 orthogroups by aligning the structures in each orthogroup to their respective *S. cerevisiae* reference structure using the *matchmaker()* function provided by *ChimeraX 1.2.5* (Goddard et al., 2018; Meng et al., 2023, 2006; Pettersen et al., 2021) using default parameters, the Needleman-Wunsch algorithm (Needleman and Wunsch, 1970) and CA-matching. UCSF ChimeraX is developed by the Resource for Biocomputing, Visualization, and Informatics at the University of California, San Francisco, with support from National Institutes of Health R01-GM129325 and the Office of Cyber Infrastructure and Computational Biology, National Institute of Allergy and Infectious Diseases. We noticed that some orthogroups had proteins that clustered together, so we refined our original sequence-based algorithms based on structural similarity.

To do this refinement, we began with US-align which is an extended version of TM-align (Zhang et al., 2022; Zhang and Skolnick, 2005) for bidirectional structure alignment. US-align was used to align each predicted protein structure to every other predicted structure that was assigned in the same orthogroup. Following this, subclusters within each of the 424 original orthogroups were generated using hierarchical clustering using *scipy.cluster.hierarchy.linkage* with default parameters. There were 12 orthogroups that contained at least one protein that did not cluster with any other protein of the orthogroup based on a linkage threshold of 0.2 (*scipy.cluster.hierarchy.dendrogram* labelled as C0 with color_threshold=0.2. This subclustering resulted in 29 orthogroups being split into two subclusters, except in the case of OG2228 containing Psd2p from *S. cerevisiae* which was split into three subclusters. This resulted in a total of 454 refined structural orthogroups uniquely identified by the orthogroup and a reference structure within that cluster.

Structure alignments for the refined structural orthogroups were then extracted as subsets of the original reference-based structure alignments and used as the basis for calculations of mapping ratios, conservation ratios and structural and metabolic analysis. For these calculations 25 refined structural orthogroups were discarded that had no *S. cerevisiae* reference structure assigned, leaving 429 refined orthogroups. From 531 remaining reference structures in *S. cer.* two structures couldn’t be assigned to any cluster and were therefore neglected for further analysis resulting in a final number of 529 reference structures.

For calculation of dN/dS as well as creating phylogenetic trees for each orthogroup, new multiple structure alignments were generated for each refined structural orthogroup without mapping to the reference structure by running US-align on the refined structural orthogroups with the option −mm 4.

For these alignments we then filtered out 135 structures whose sequences were less than 80% of the median length of their structural orthogroup, and then filtered out 22 structural orthogroups that had 3 or fewer sequences. Each refined orthogroup was then assigned a reference structure for naming purposes, which was from *S. cerevisiae* in all but 7 of the refined orthogroups. This left 432 refined orthogroups that were carried forward for further analysis. Genes were designated as paralogs if they were from the same species in the same refined orthogroup.

### Multiple sequence alignments and phylogenetic trees

Multiple sequence alignments (MSAs) were generated for the 432 refined orthogroups for which short sequences and orthogroups with 3 or fewer structures were removed. Structure informed alignments (US-Align) versions were generated with the US-Align program with the option −mm4. To generate phylogenetic trees for each MSA alignment were first trimmed using clipkit version 1.4.1 with default parameters (Steenwyk et al., 2020). IQtree version 2.1.4-beta was then run on each alignment using default parameters that allow different evolutionary models to be identified and used for each sequence. The options –bb 1000 and -alrt 1000 to evaluate the quality of each tree (Minh et al., 2020).

### Metabolic network visualisation

Visualisation of the yeast metabolic pathways was performed using iPath3 (Darzi et al., 2018). Of the 529 *S. cerevisiae* structures from 429 orthogroups we analysed, 448 were present in the iPath3 database metabolic network, covering 62.5% of the 717 *S. cerevisiae* uniprot IDs present in the iPath3 metabolic network. Overall, the iPath3 metabolic network contained 243 more proteins than the yeast pathways database which we used to identify enzymes and orthogroups of interest. Colour and width of each element in the metabolic map was determined by the average conservation scores for the proteins assigned to that element in iPath 3 for the full KEGG metabolic map (01100) subsetting on *S. cerevisiae* (sce).

### Phenotype selection

Growth phenotypes in 72 different phenotypes for various budding yeast species was taken from (Lu et al., 2021), which assembled data originally collected in (Kurtzman et al., 2011). 6 conditions which did not vary across our selected species were removed, and 44 conditions which had data from fewer than 5 species that can grow and fewer than 5 species that can not grow under the respective condition were removed.

### Protein abundance measurement

#### Cultivation

Strains were streaked on CN1 2% agar plates and incubated at 25°C for 48 h. Single colonies were picked, inoculated into 60 ml CN1 liquid medium (300 ml flask total volume) and incubated for 24 h at 25°C with shaking at 200 r.p.m. (Sartorius Certomat IS Shaker). Subsequently, optical density at 600 nm (OD600) was recorded and 20 ml of culture were centrifuged at 1,500 x g for 5 minutes at RT (Eppendorf Centrifuge 5810 R). After centrifugation, the supernatant was discarded, and the cell pellet was resuspended in 20 ml of CN1 fresh medium (flask size was adjusted to 100 ml to keep the same liquid-to-air ratio of pre-cultures). Cultures were placed back into the incubator for shaking with the same settings (25°C, 200 r.p.m.). After 6 h of incubation, corresponding to mid-log phase,2 ml of culture were sampled into screw cap tubes. A small volume of 50 μl was removed for OD600 measurement while the remaining sample was immediately centrifuged at 21,000g (Thermo Scientific Heraeus Fresco 21 Microcentrifuge) for 1 minute at 4°C, then the supernatant was carefully removed by inversion and the cell pellet was flash frozen with liquid nitrogen. Finally, ∼130 μg of pre-aliquoted glass beads (425-600 μm) were added to each sample tube with a small funnel and frozen at −80°C until further processing.

Processing QC samples were generated by cultivating the lab strain *S. cerevisiae* BY4741ki in rich Yeast Peptone Dextrose (YPD) medium. A single flask of 400 ml YPD was inoculated with BY4741ki directly from the cryostock. The culture was incubated for 20 h at 30°C with shaking at 200 r.p.m. (Sartorius Certomat IS Shaker), then distributed into aliquots of 4 x 1e+08 cells. Cells were harvested by centrifugation at 21,000g (Thermo Scientific Heraeus Fresco 21 Microcentrifuge) for 1 minute at 4°C, then the supernatant was removed, and the cell pellet was frozen at −80°C until further processing.

#### Cell lysis

Frozen pellets were topped with 200 μl of lysis buffer (7M urea, 0.1M ammonium bicarbonate) and lysed in a bead-beater (SPEX, SamplePrep 1600 MiniG) for 5 minutes at maximum speed (1,500 r.p.m.). The procedure was performed three times for a total of 15 minutes of beating per sample and each cycle was followed by 5 minutes of cooling down on ice to avoid sample overheating.

After 1 min of centrifugation at 15,000 r.p.m. (Thermo Scientific Heraeus Fresco 21 Microcentrifuge) at 4°C to clear the lysate, protein concentration was measured by Pierce 660nm Protein Assay (ThermoFisher). The volume corresponding to 100 μg protein was transferred to a 96-well 2 ml deep-well plate (Eppendorf) and total volume was adjusted to 200 μl with lysis buffer.

#### Sample preparation for mass spectrometry

Sample preparation was adapted from Messner C.B., Demichev V., Bloomfield N. et al., Nat Biotechnol., 2021, 39, 846–854, doi: 10.1038/s41587-021-00860-4). Samples were treated by addition of 20 μl of 55-mM DL-dithiothreitol (final concentration 5 mM). The plate was mixed for 2 minutes at 1,000 r.p.m., and incubated for 1 h at 30 °C. After incubation, samples were cooled on ice for 5 minutes. Subsequently, 20 μl of 120 mM iodoacetamide was added (final concentration 10 mM), then mixed for 2 minutes at 1,000 r.p.m., and incubated for 30 min in the dark at room temperature. Subsequently, 460μl of 100-mM ammonium bicarbonate was added. Samples were mixed for 90 seconds at 1,000 r.p.m., and an aliquot of 500 μl was transferred to pre-filled trypsin/LysC plates (Waters, 96-well plate 700 μl round, 4 μg of trypsin/LysC). After incubation of the samples for 17 h at 37 °C with shaking at 750 r.p.m. (Benchmark Scientific, Incu-Mixer Microplate vortexer), trypsin/LysC were deactivated by addition of 17 μl of 30% formic acid (final concentration 1%). The digestion mixtures were cleaned using C18 96-well plates (96-Well BioPureSPN, PROTO 300 C18, 35-350µg max capacity, The Nest Group, no. HNS S18V-L). For solid-phase extraction, 1-min centrifugation steps at the described speeds (Eppendorf Centrifuge 5810 R) were applied to force liquids through the stationary phase. A liquid handler (Beckmann Coulter Biomek i7) was used to pipette the liquids onto the material. The plates were conditioned with methanol (200 μl, centrifuged at 50g), washed twice with 50% acetonitrile (ACN, 200 μl, centrifuged at 150g and flow-through discarded) and equilibrated three times with 0.1% formic acid (200 μl, centrifuged at 150g, respectively, and flow-through discarded). Then, 500 μl of digested samples was loaded (centrifuged at 550g) and washed two times with 0.1% formic acid (200 μl, centrifuged at 150g). After the last washing step, the plates were centrifuged once more at 200g before elution of peptides in three steps, each with 110 μl of 50% ACN (200 g), into a collection plate (Waters, 96-well plate 700 μl, round). The collected material was completely dried on a vacuum concentrator (Thermo Scientific, Savant SpeedVac SPD300) and redissolved in 90 μl 0.1% formic acid by mixing for 5 minutes at 1,000 rpm before transfer to a 96-well plate (700 μl round, Waters, no. 186005837). Peptide concentration was measured using Pierce Quantitative Peptide Assay (ThermoFisher Scientific). All shaking was performed with a thermomixer (Eppendorf Thermomixer C) and, for incubation, a Benchmark Scientific, Incu-Mixer Microplate vortexer was used.

#### Liquid chromatography mass spectrometry

Samples were analysed on a Bruker timsTOF Pro mass spectrometer, coupled to a Dionex Ultimate 3000 µsystem (Thermo Fisher Scientific). Prior to MS analysis, 1µg peptides were chromatographically separated with a 30 min gradient on a Waters HSS T3 column (300um x 150mm, 1.8um) heated to 40°C, using a flow rate of 5 µl /min where mobile phase A & B are 0.1% formic acid in water and 0.1% formic acid in ACN, respectively. The active gradient increases from 2% to 40% B in 30min.

For diaPASEF acquisition, the electrospray source (Bruker Apollo II source, Bruker Daltonics) was operated at 4500 V of capillary voltage, 5.0 l/min of drying gas and 200 C° drying temperature The dia-PASEF windows scheme was as followed: we sampled an ion mobility range from 1/K0 = 0.6 to 1.60 Vs/cm2 using equal ion accumulation and ramp times in the dual TIMS analyzer of 100 ms, each cycle times of 0.5 s. The collision energy was lowered as a function of increasing ion mobility from 59 eV at 1/K0 = 1.6 Vs/cm2 to 20 eV at 1/K0 = 0.6 Vs/cm2. For all experiments, TIMS elution voltages were calibrated linearly to obtain the reduced ion mobility coefficients (1/K0) using three Agilent ESI-L Tuning Mix ions (m/z, 1/K0: 622.0289, 0.9848 Vs/cm2; 922.0097, 1.1895 Vs/cm2; and 1221.9906, 1.3820 Vs/cm2).

#### Data processing

Spectra deconvolution, protein identification and relative quantification was performed using DIA-NN 1.8 (Data-Independent Acquisition by Neural Networks) [Demichev V. et al., Nat Methods. 2020; 17(1): 41–44, doi: 10.1038/s41592-019-0638-x]. Peptide search was performed in library-free mode, meaning that deep learning was used to generate a new *in silico* spectral library from the Uniprot reference proteome available for each species (downloaded on Sept 13th, 2021). The following settings were applied: output was filtered at 0.01 FDR, minimum fragment m/z was set to 200, maximum fragment m/z was set to 1800, N-terminal methionine excision was enabled, *in silico* digest involved cuts at K*,R*, maximum number of missed cleavages was set to 1, minimum peptide length was set to 7, maximum peptide length was set to 30, minimum precursor m/z was set to 350, maximum precursor m/z was set to 1300, minimum precursor charge was set to 1, maximum precursor charge was set to 4, cysteine carbamidomethylation was enabled as a fixed modification, scan window radius was set to 10, thread number was set to 40, mass accuracy was fixed to 1e-05 (MS2) and 1e-05 (MS1). Finally, data were processed in R by filtering proteotypic precursors at sample fraction 80% within all samples of the same species, and a minimum of 3 precursors were used for protein quantification. The minimum threshold for Global.Q.Value, Global.PG.Q.Value, Q.Value, PG.Q.Value was set to 0.01.

### Mapping structures within an orthogroup

To generate a one-on-one assignment of the amino acids between different structures within an orthogroup, we utilised the structures that were aligned to the reference structure of *S. cer..* In the first step we spanned a tree based on the euclidean distances between the C_ɑ_-atoms of the reference structure and the C_ɑ_-atoms of the aligned structures. Using a cut-off of 2 Å, we generated a C_ɑ_ mapping matrix for each orthogroup. Each amino acid that could not be mapped to the reference structure was excluded from later analysis. An example of the mapping is shown Figure S1.3D. Based on this mapping matrix, structural information as described below were projected onto the alignments. To calculate the mapping rate, the portion of residues that could be mapped to the reference structure versus the length of the reference structure was calculated. The mapped length was calculated as the number of amino acids of the original protein chain that could be mapped to the reference structure. The conservation ratio denotes the portion of no amino acid changes with respect to the corresponding reference structure divided by the length of the mapped parts. Overall analysed orthogroups a median CR of 62.9% (IQR: [53.6%, 71.2%], total: [24.5% to 89.2%]) was observed and the range was 24.5% to 89.2% (10% quantile: 47.8%, 90% quantile: 77.2%).

### Binding site extraction

Binding sites were defined in two different ways. For the direct coordination sphere the binding site (and for the enrichment analysis also the active sites) were downloaded from UniProt in November 2022 (The UniProt Consortium, 2023). To account for the physicochemical environment, we used crystal structures extracted from RCSB Protein Data Bank (RSCB.org) (Berman et al., 2000) and the crystal structure annotation of Yeastmine (http://yeastmine.yeastgenome.org/ accessed 7 Oct, 2021) (Wong et al., 2023). Each structure was aligned to the corresponding reference structure and all hetero-atoms were extracted. To remove crystalizing agents, we removed halides and small anions like nitrate, azide, sulphate and phosphate as well as alkali metals, small cations like ammonium and heavy metals. In addition, water molecules were removed as well. All other heteroatoms were treated as ligands. To determine the binding site and its physicochemical environment, we selected all amino acids where at least one atom was within a distance of 5Å to bound ligands or cofactors from X-ray crystallographic and cryo-EM structures from *S. cerevisiae* in our structural alignments.

### Structural information extraction

#### Estimating properties and enzyme cost

All properties were estimated on the re-clustered orthogroups. Thus, orthogroups containing more than one cluster were split. In addition, the analysis was performed for every single reference structure in *S. cer.*.

The solvent accessible surface area (SASA) was evaluated using the Shrake-Rupley algorithm (Shrake and Rupley, 1973), the summarised secondary structure estimates were determined using the DSSP algorithm (Kabsch and Sander, 1983) as implemented in *MDTraj* 1.9.7 (McGibbon et al., 2015). The pLDDT values were extracted from the predicted AlphaFold2 structures. For the relative amino acid content, the ratio between a specific amino acid in the structure and the overall length of the protein chain was calculated. Thus, the relative amino acid content per structure over all amino acids sums up to one. To take into account other physicochemical properties we also included the amino acid hydrophobicity (https://www.genome.jp/entry/aaindex:CIDH920105) (Cid et al., 1992) and other amino acid specific properties like the isoelectric point or the amino acid polarity (Gao et al., 2023) of the whole protein chain.

For estimating the protein costs, we included several different cost terms: Molecular weight, energy equivalents (Akashi and Gojobori, 2002; Craig and Weber, 1998; Wagner, 2005), number of reaction steps(Craig and Weber, 1998), nutrient usage in the context of metabolic flux models (FBA) (Barton et al., 2010), and costs of generating the enzymes required to synthesise the protein (Chen and Nielsen, 2022). For calculating an averaged normalised cost, each cost metric was bound between 0 and 1 and the mean per amino acid was calculated.

Properties were estimated for the whole non-aligned structures, the reference structure only and certain extracted structural features based on the amino acids that could be mapped to the reference structure (see, following subsection). For the calculation of the average properties per orthogroup only the structures belonging to the cluster of the reference structure as described above were used.

#### Extracted Structural features

To utilise one advantage of our structural predictions, we calculated most of the estimated properties not only for the mapped parts, but also for specific structural features. For this the mapped structures were divided into groups using different subset. For the secondary structural elements, we used the summarised DSSP information to distinguish helical, extended and random coil structural elements. To distinguish the surface and core residues we calculated the maximal SASA per amino acid for non-terminal amino acids. For this we constructed an artificial helical protein chain GX1GX2G[…]GX20G with torsion angles \phi = 50° and \psi = 45° to achieve the maximal surface accessibility per amino acid (Tien et al., 2013). Core residues were defined as buried residues with a maximal relative accessible surface area of 25% (Gong et al., 2017; Momen-Roknabadi et al., 2008). For the binding sites the definitions as given above were used. Also, the crystal structure exclusive part of the binding site that was not defined in UniProt was extracted. In addition, the active side definition extracted from UniProt was included. Furthermore, positions of reference structure matching, fully conservation as well as positions with at least one amino acid or amino acid type mutation were tested. Another subset for amino acids with an pLDDT greater than 0.7 was estimated. As a last selection the intracellular location (Weill et al., 2019) was included splitting into membrane-bound, cytosolic, luminal/extra-cellular and signalling as well as into a combination of cytosolic and luminal/extra-cellular.

#### Grouping of amino acids

For the grouping of amino acids, we decided into 7 different groups. The 5 main groups were divided by physicochemical properties: (I) Acidic amino acids (D, E), basic amino acids (H, K, R), polar amino acids (C, N, Q, S, T), apolar amino acids (A, I, L, M, V) and aromatic amino acids (F, W, Y). The canonical amino acids G and P were treated as separate groups due to their unique backbone dynamics. In the case of G (flexible) a higher backbone flexibility can be achieved due to the missing side chain. In the case of P (break) a decreased backbone flexibility as well as influence on neighbouring amino acids are observed due to the ring formation of the side chain with the amino acid backbone.

### Evaluated experimental data sets and databases

To investigate the relationship of structural evolution and experimental observables, we investigated several experimental data sets. In addition, we included different databases for system level analysis. A summary of the used sources is found in the following table.

### Flux and kcat calculations

For flux calculation genome-scale metabolic models for 329 yeast species were utilised (Lu et al., 2021). A total of 66 conditions, encompassing various carbon sources combined with minimal media, were employed as inputs for the simulations. The maximum uptake rate for carbon sources was defined by setting the lower bound at −1 Cmmol/gDW/h. Growth was selected as the objective function for optimization. For each orthogroup, the mean fluxes across all conditions, excluding zero, were calculated to determine the average flux for each OG in each species. Due to the logarithmic transformation 0s were excluded for the calculation of the median flux. For the calculation of the number of species with Flux > 0, all 329 species were included.

For the estimation of *kcat* values, we utilised species-specific models for various yeast species. Each model was meticulously refined, with reversible enzymatic reactions split into forward and backward processes and reactions catalysed by isoenzymes segmented into separate reactions with individual enzyme complexes. SMILES information for substrates were extracted by mapping substrates into MetaNetX database (Moretti et al., 2021). Enzyme information, extracted from the model’s gene-reaction rules (grRules), was complemented by protein sequences obtained via protein IDs from species-specific protein FASTA files. This dataset, encompassing reaction IDs, substrate names, SMILES information, and protein IDs, formed the basis for our deep learning kcat prediction model, DLKcat. Entries with non-specific SMILES or asterisks (*) in protein sequences were excluded from prediction. Since each enzyme can utilise multiple substrates in multiple reactions, we first evaluated summarization within the reactions followed by the reactions per enzyme itself by different methods (maximal value, minimal value, mean or median). On orthogroup level in addition the standard variation as well as the coefficient of variation were calculated. In the course of the manuscript, we only report the (within and between reactions per enzyme) median summarised log2-transformed values.

### Enrichment analysis

For categorical data collections such as gene locus, enzyme class, metabolic pathway or GO slim terms we performed an enrichment if not mentioned otherwise of the top 25 % as well as lowest 25 % of the analysed observables. To get a more detailed understanding we analysed enzyme classes on the first level as well as combinations of the first and the second or third level respectively. For the enrichment analysis we performed a Fisher’s exact test as implemented in *scipy 1.8.1* (Virtanen et al., 2020). Within each collection, we performed Benjamini-Hochberg multiple testing correction (Benjamini and Hochberg, 1995) as implemented in *statsmodels 0.13.2* (Seabold and Perktold, 2010). The significance level was set to adj. p-value < 0.05. To account for the analysed protein space, we have only included the analysed orthogroups as a background truth. In addition, we calculated the area under the receiver operating characteristic curve (abbrev. AUC) as implemented in *scikit-learn 1.1.1* (Pedregosa et al., 2011). For the calculation of the enrichment factor, we divided the number of observations by the number of expected observations assuming a random distribution.

### Correlation and statistical analysis

For statistical testing as well as correlation calculations *scipy 1.8.1* (Virtanen et al., 2020) was used. Depending on the analysis we distinguished between the linear Pearson correlation coefficient *r*, the rank-order Spearman correlation coefficient ⍴ (Spearman, 1904) or for discrete data Kendall’s τ coefficient (Kendall, 1945, 1938) using *variant*=*”b”*. To account for multiple testing, the data set was split into data extracted from the structural alignment and experimental data. Benjamini-Hochberg multiple testing correction (Benjamini and Hochberg, 1995) was applied as implemented in *statsmodels 0.13.2* (Seabold and Perktold, 2010) for all calculated p-values within the same data, the same correlation measure and the same property (noted on the x-axis of the respective plots). For statistical testing we used the nonparametric Wilcoxon-Mann-Whitney test (Mann and Whitney, 1947; Wilcoxon, 1945) for non-paired samples as well as the nonparametric Wilcoxon signed-rank test (Wilcoxon, 1945) for paired samples. For comparing two continuous distributions we used the two-sided, two-sample Kolmogorov-Smirnov test (Smirnov, 1948). In all cases missing values were omitted. For the visualisation of distributions a smoothing function with a smoothing-factor ≤ 1E-8 was applied, which might cause artefacts at the borders of the distributions.

### Network clustering

To extract regions that never changed during evolution, within each orthogroup, we first identified all fully conserved residues in the alignment. In a next step we set up a network of these residues using *igraph 0.9.9* (Csárdi and Nepusz, 2006) in which nodes are connected if the residue’s C_ɑ_-atoms are not more than 10 Å apart. Using a fixed seed of 42, we clustered the network using the Leiden algorithm (Traag et al., 2019) as implemented in *leidenalg 0.8.9* using *CPMVertexPartition* and a resolution parameter of 0.05. The clusters were extracted and plotted using *igraph*. To remove artefacts, the minimum number of amino acids per cluster was set to 5.100.

### Structural representation

For the evaluation of the positions of the positive selected residues, we centred the CIII-CIV-complex and rotated it onto the principal axis using *GROMACS 2021.5* (Abraham et al., 2015). We extracted the y-coordinates as they resemble the orientation of the membrane-inserted complex. For structural visualisations we used *VMD 1.9.4*. *VMD* is developed with NIH support by the Theoretical and Computational Biophysics group at the Beckman Institute, University of Illinois at Urbana-Champaign.

### Evolution rate (dN/dS) calculations

For evolution rate (dN/dS) calculations, nucleotide alignments were required. Nucleotide sequences were obtained from either model organism databases (*C. albicans* and *S. cerevisiae*), NCBI’s datasets resource [https://www.ncbi.nlm.nih.gov/datasets/], NCBI’s genome database by downloading data from individual contigs, or (Shen et al., 2018). These sequences were then threaded onto protein alignments using *phykit v1.11.15* (Steenwyk et al., 2021). The alignments were then strictly trimmed using *clipkit v1.4.1* with options *-m gappy -g clipkit* (Steenwyk et al., 2020). Three refined orthogroups for which the strict trimming was less than 25% of the median sequence length were removed at this stage, OG1306_REF_Scer_AF-P38298-F1-model_v2, OG1746_REF_Scer_AF-P32642-F1-model_v2, and OG2228_geotrichum_candidum OG2228 43_6293. Strictly trimmed alignments, and phylogenetic trees when necessary, were then formatted and sequences given shorter names in order to work with CodeML.

Two strategies were used to estimate average dN/dS values for entire orthogroups. The Yang and Nielsen method (YN00) method (Yang and Nielsen, 2000) and the one-ratio model (M0) (Goldman and Yang, 1994; Yang and Nielsen, 1998), which is informed by phylogenetic trees. Values from the one-ratio model were used for enrichment calculations, and a filter with dS>3 was applied as described in the main text.

To identify dN/dS estimates at the amino acid level for example orthogroups containing Pox1p and proteins from the ETC, we followed a two-stage process using the *codeml* function from *PAML v4.10.6* (Yang, 2007). First, we used the free ratio branch model (M1) (Yang, 1998; Yang and Nielsen, 1998). Any branches for which dN/dS was greater than 1.1 in that test, were then tested using the branch-site test for positive selection (BS) (Yang et al., 2005; Zhang et al., 2005). If the χ2 threshold between model A and model A with ω fixed at 1 was greater than 3.84, then residues for which the Bayes Empirical Bayes posterior probability of dN/dS>1 was greater than 0.95 were considered under positive selection.

## Acknowledgements

We would like to thank Jurg Bahler, Jason Yu, Julia Muenzner, Johannes Hartl, Chris Jakobsen, Tauqeer Alam, Matt White, Sandra Alvarez-Carretero and Ziheng Yang for scientific advice. In addition, we would like to acknowledge Jessica Tamanini for editing support and Lukasz Szyrwiel, Daniela Ludwig and Michael Mülleder for preparation and support for the proteomics measurements. OL and NC are supported by the European Research Council (ERC) under grant agreement ERC-SyG-2020 951475. BMH and SA are supported by Wellcome Trust grant (IA 200829/Z/16/Z). The computations for structure predictions were enabled by the Berzelius resource provided by the Knut and Alice Wallenberg Foundation at the National Supercomputer Centre. Initial computations and data storage was enabled by resources provided by Chalmers e-Commons at Chalmers. JLS is a Howard Hughes Medical Institute Awardee of the Life Sciences Research Foundation. Research in AR’s lab is supported by the National Science Foundation (DEB-2110404), the National Institutes of Health/National Institute of Allergy and Infectious Diseases (R01 AI153356), and the Burroughs Wellcome Fund.

## Author’s contribution

OL, BMH, TIG and MR conceptualised the work. OL, BMH, SV, NC, JLS and TIG developed the methodology. OL, BMH, SV, NC, JLS, LS and FL implemented the software for the analysis. OL, BMH, SV, FL, FA and CTL validated the data. OL, BMH, NC, JLS and LS performed the formal analysis and investigation of the data. SV generated the structures. AZ and MR provided resources. OL, BMH, SV, NC, JLS, LS, FL, FA and SKA were involved in data curation. OL, BMH and MR wrote the original draft. OL and BMH prepared the figures. TIG and MR supervised and OL, BMH and MR administered the project. JN, AR, AZ and MR provided funding or resources to at least one coauthor.

## Code Availability Statement

Functions used to map the structures and visualise the data are uploaded to https://github.com/OliverLemke/structure_comparison and https://github.com/OliverLemke/basic_plots. Functions used to assemble MSAs, phylogenetic trees and do evolutionary selection analysis are uploaded to https://github.com/heineike02/diverse_yeast. Most code used for generation of data and plots will be available upon publication as a supplement. Missing code will be available upon request.

## Data Availability Statement

Proteomics data will be uploaded to the PRIDE database prior to publication. All other data will be uploaded to figshare and made available upon publication. Supplemental tables are listed below:

### Supplemental Tables

| Table | Notes |
| --- | --- |
| S1.1 | Lists of selected species and source of sequences for each species. |
| S1.2 | List of structures that failed to calculate. |
| S1.3 | Average pLDDT and percentage of structures calculated for each orthogroup. |
| S1.4 | Orthogroup, species, UniProt ID, Length, Cluster and pLDDT for structures that we use for analysis. Highlight if structure was predicted during the course of the study or extracted from AlphaFold DB. |
| S1.5 | Table of refined clusters generated from original orthogroups based on TM-align scores. Includes list of *S.cerevisiae* genes in each cluster. |
| S1.6 | Selected metrics and calculated values for each structural orthogroups, like CR, MR, Mapped length, Amino acid content and protein abundances. |
| S5.1 | General descriptions on the network clusters. To remove artefacts, clusters have to have at least a size of 5. The median value as well as the interquartile range (IQR) are presented for both binding site definitions. |
| S6.1 | Key residues highlighted in Figure 6C for the orthogroup containing *S. cerevisiae* Pox1p across different phylogenetic orders. |

---

**Figure S1.1:**
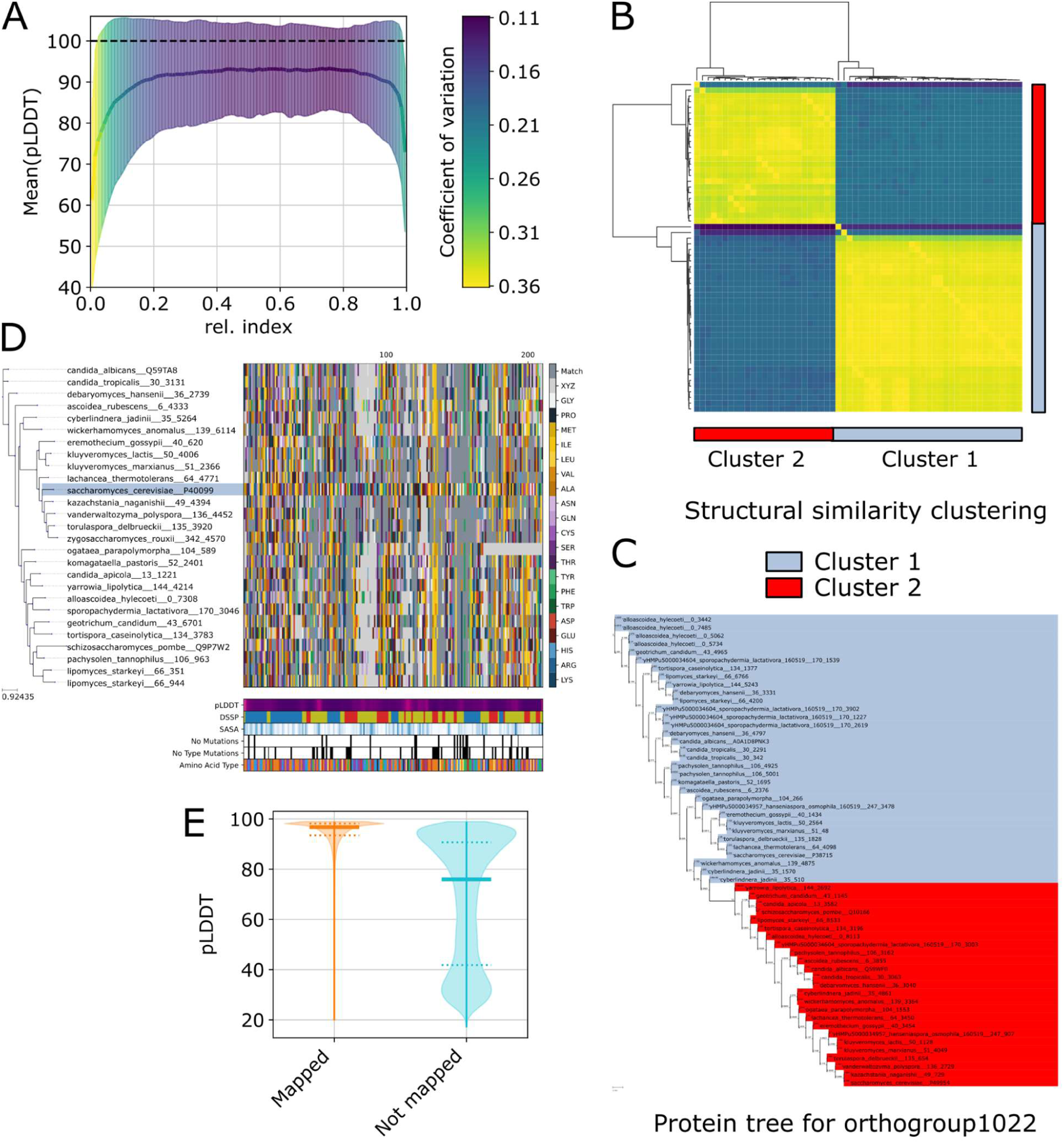
The predicted protein structures are well-structured, but small orthogroup refinements are required. A) Mean pLDDT score as calculated by Alphafold2 versus the relative, length-normalised amino acid index for all our simulated structures, divided into 100 bins, due to varying protein lengths. Error bars denote the standard deviation coloured by the coefficient of variation. B+C) Structural alignment-based orthogroup refinement illustrated on orthogroup 1022. B) Hierarchical clustering of structures using US-align scores. The two resulting clusters are highlighted. Cluster 1 (blue) contains the reference structure *S. cerevisiae* Gre3p (Uniprot: P38715) and is named OG1022_REF_Scer_AF-P38715-F1-model and cluster 2 (red) contains Nit3p (Uniprot: P49954) and is named OG1022_REF_Scer_AF-P49954-F1-model. C) Phylogenetic tree based on US-align structural alignment of the entire original orthogroup 1022. The clusters based on hierarchical clustering correspond to structures that cluster together in the protein tree. D) Example of the structural alignment with respect to the reference structure. In addition, in the bottom panel other features, like pLDDT, SASA, DSSP, fully conserved residues and fully conserved amino acid types are depicted. Colour according to amino acids. In case of light grey, no residue could be aligned to the reference. A dark grey colour indicates an agreement with respect to the reference structure for the respective amino acid. The more dark grey, the higher the conservation ratio. E) Maximum-normalised violin plot of the pLDDT for the density distribution containing every single residue grouped by residues that could be mapped (n=5,527,262) and that could not be mapped (n=1,454,754) respectively.

**Figure S2.1:**
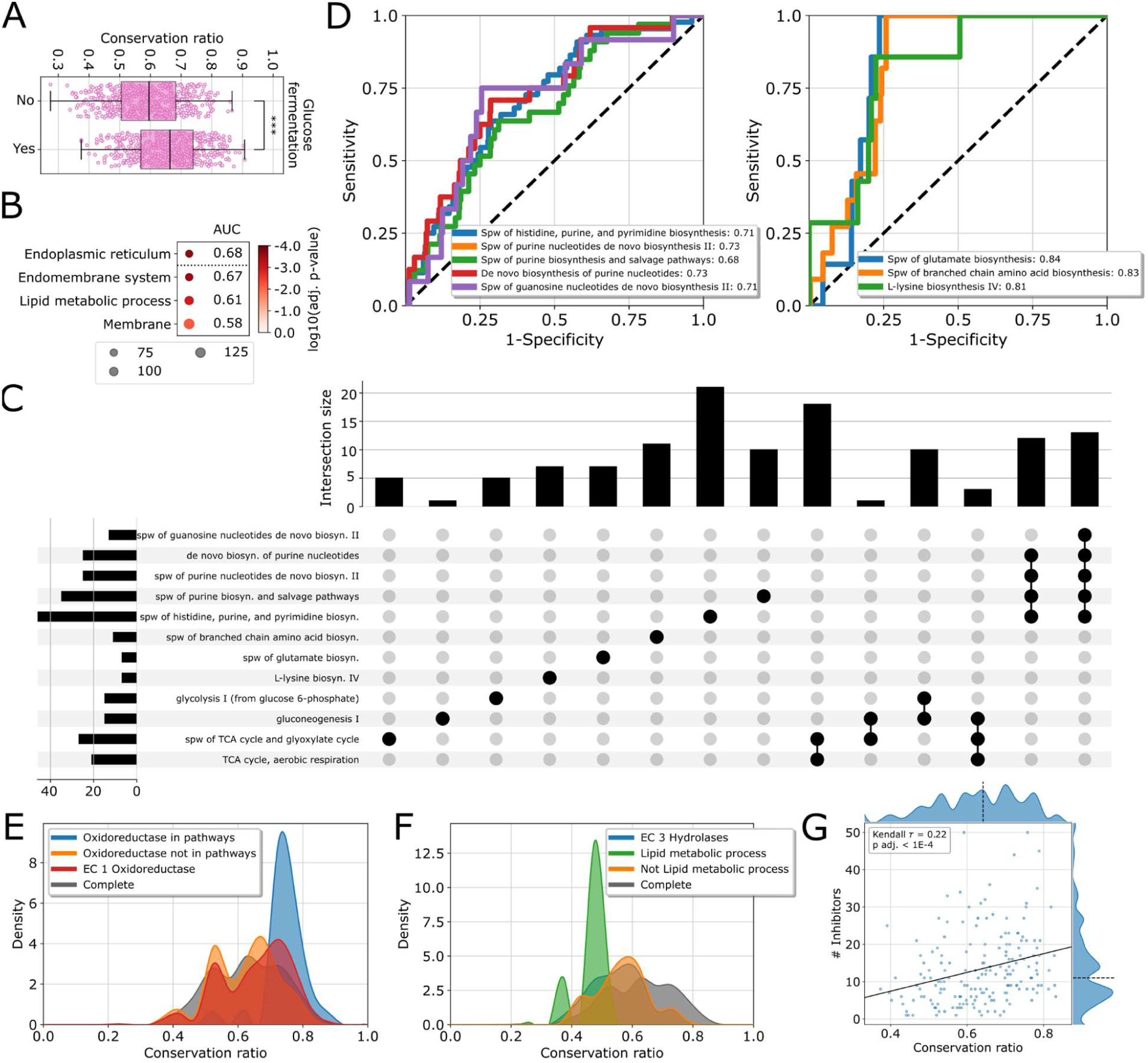
Metabolic factors constraining structural evolution in *Saccharomycotina.* A) Boxplot of the conservation ratio for species that can grow utilising glucose fermentation versus non-fermenting species. *** indicates a p-value < 1e-4, Wilcoxon signed-rank test. B) Significant GO slim enrichments of the first quartile of the orthogroups containing proteins that show the largest differences in mean conservation ratio between species that ferment and species that do not ferment glucose. Size indicates the number of genes associated with the GO slim term in our selection, and colour indicates adjusted p-value. In addition, the area under the receiver operating characteristic curve (abbrev. AUC) is shown. C) Upset-plot of the significantly enriched pathways, ordered by group dependency. D) ROC-curve for highly enriched pathways. The number in the legend indicates the AUC. The dashed line denotes the identity line representing random sampling. E) Distribution of the mean conservation ratio of orthogroups assigned to oxidoreductases (red). In addition, the distributions for oxidoreductases that are (blue) or that are not members of the enriched pathways from (D) and Figure 2D (orange) are depicted. The average distribution for all orthogroups is shown in black. F) Distribution of the mean conservation ratio of orthogroups assigned to hydrolases (blue). In addition, the distributions for hydrolases that are (green) or that are not (orange) members of the term “lipid metabolic process” are depicted. The average distribution for all orthogroups is shown in black. G) Mean conservation ratio versus the number of inhibitors. Solid line is the best linear fit and the dashed line denotes the axis median.

**Figure S3.1:**
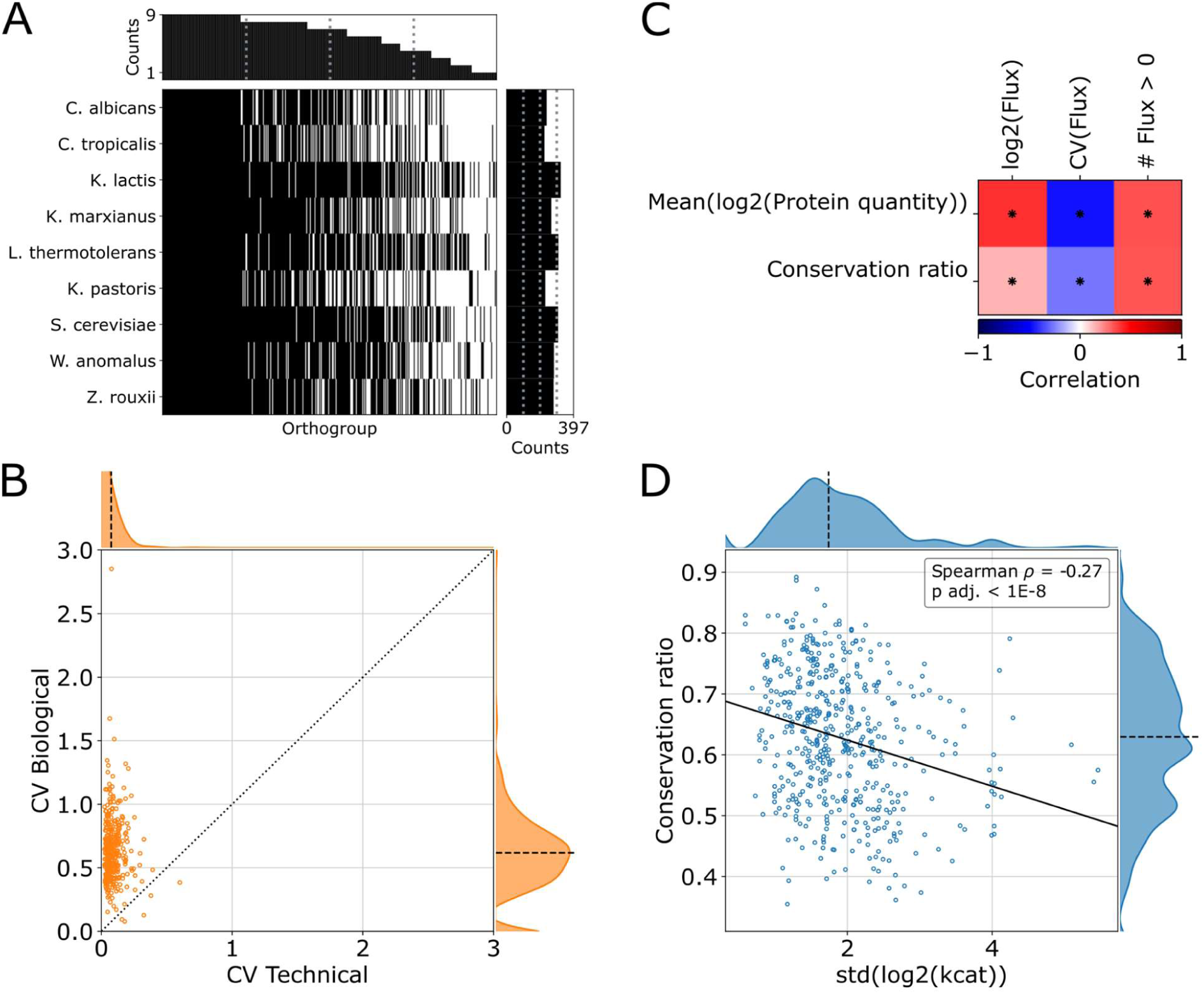
The proteomics data are of high-quality. Metabolic properties influence structural evolution. A) Completeness of proteomic measurements for the selected orthogroups. For 397 of the original orthogroups protein abundance was measured in at least one species. A black line indicates a measured protein abundance of the orthogroup in the respected species. The bar plots account for the row and column sum. Grey dotted lines indicate the first, second and third quartiles. B) Scatter plot of the technical coefficient of variation (quality control samples) versus the biological coefficient of variation (measured species). The dotted line denotes the identity line and the dashed line denotes the axis median. C) Spearman correlation of log2-transformed median predicted flux, coefficient of variations and Kendall’s τ correlation of the number of budding yeast species (n=329) in which a flux>0 was predicted (Lu et al., 2021) versus mean protein abundance and mean orthogroup conservation ratio. A * indicates an adj. p-value < 0.05. D) Mean conservation ratio versus standard deviation of the log2-transformed averaged *kcat*. Solid line is the best linear fit and the dashed line denotes the axis median.

**Figure S4.1:**
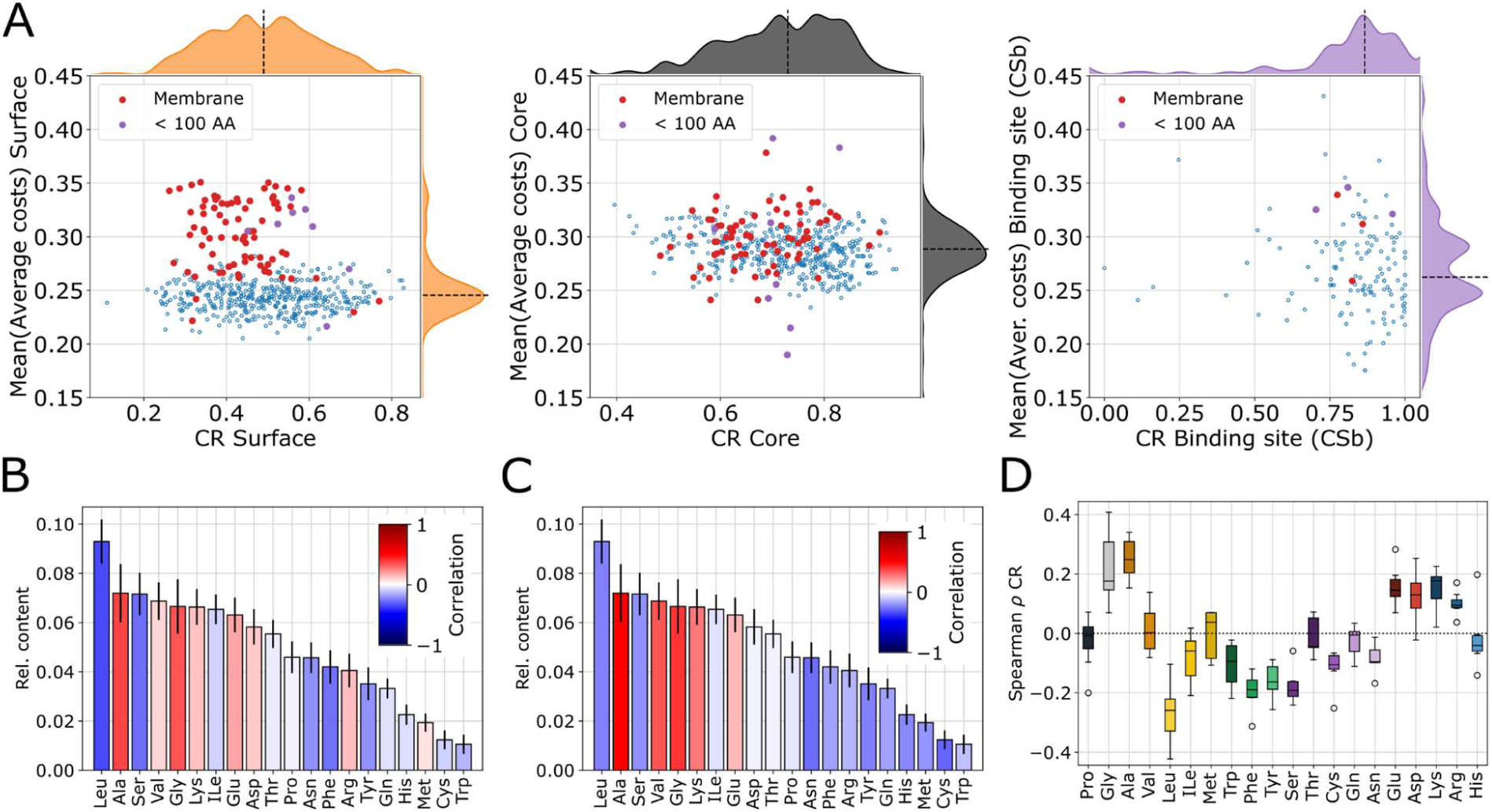
Amino acid costs are optimised depending on the structural feature and the physicochemical environment. A) Conservation ratio of the surface (left), core (centre) and binding site (CSb) (right) versus their respective mean normalised costs. Membrane bound (red) and short protein chains (purple) are highlighted respectively. For the surface a bimodal cost distribution is observed. The dashed line denotes the axis median. B+C) Bar plot of the median relative amino acid content sorted in descending order. The error bars denote the MAD. The colouring of the bars indicates the Spearman correlation of the relative amino acid content of the entire protein versus B) the mean overall conservation rate and C) the log-transformed protein abundance. D) Boxplot of the Spearman correlation coefficients as shown in Figure 4E (conservation ratio vs. relative amino acid content of different structural features). The boxes denote the first and third quartile, the whiskers in maximum 1.5-times the interquartile range. Outliers are shown as single dots.

**Figure S5.1:**
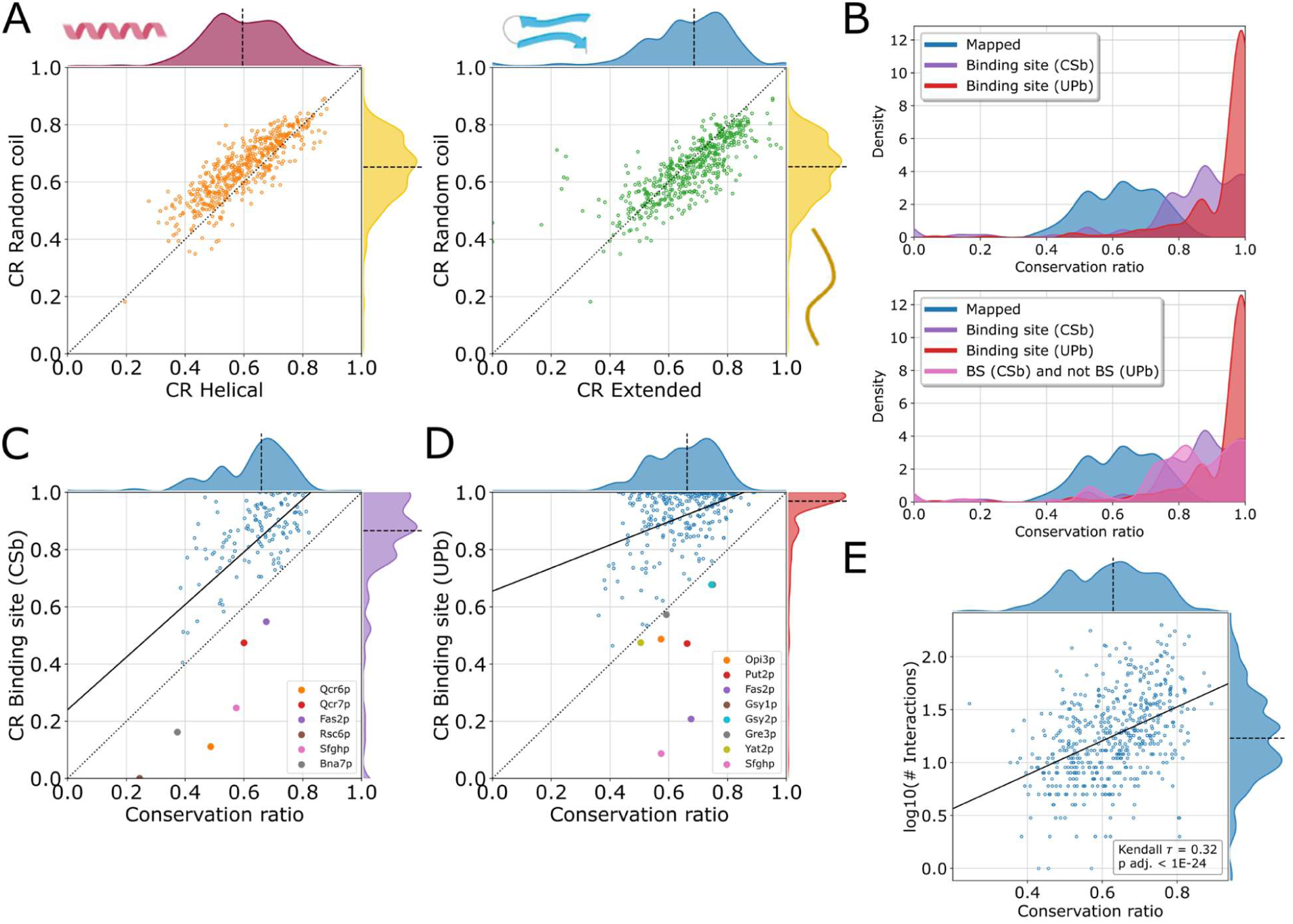
There is a hierarchy of evolution of structural features with surface residues being least constrained and binding sites the most. A) Mean conservation ratio of the random coiled versus mean conservation ratio of the helical parts (left) and the extended parts (right) respectively. Dashed line indicates the identity line and the dashed line denotes the axis median. B) Distribution of the mean conservation ratio for all mapped parts (blue) and only for the binding site (purple and red). The purple distribution depicts the CSb definition, the red distribution the UPb definition (top). In the lower panel the pink distribution accounts for residues present in the purple distribution (CSb) and not present in the red one (UPb). C+D) Mean conservation ratio of the overall protein versus the conservation of the respective binding sites for C) the CSb definition and D) the UPb definition. Orthogroups below the dashed identity line are highlighted. Solid line is the best linear fit. E) Mean conservation ratio versus the log10-transformed number of unique physical interactions taken from BioGRID (Oughtred et al., 2021) via the Yeastmine interface (Wong et al., 2023).

**Figure S6.1:**
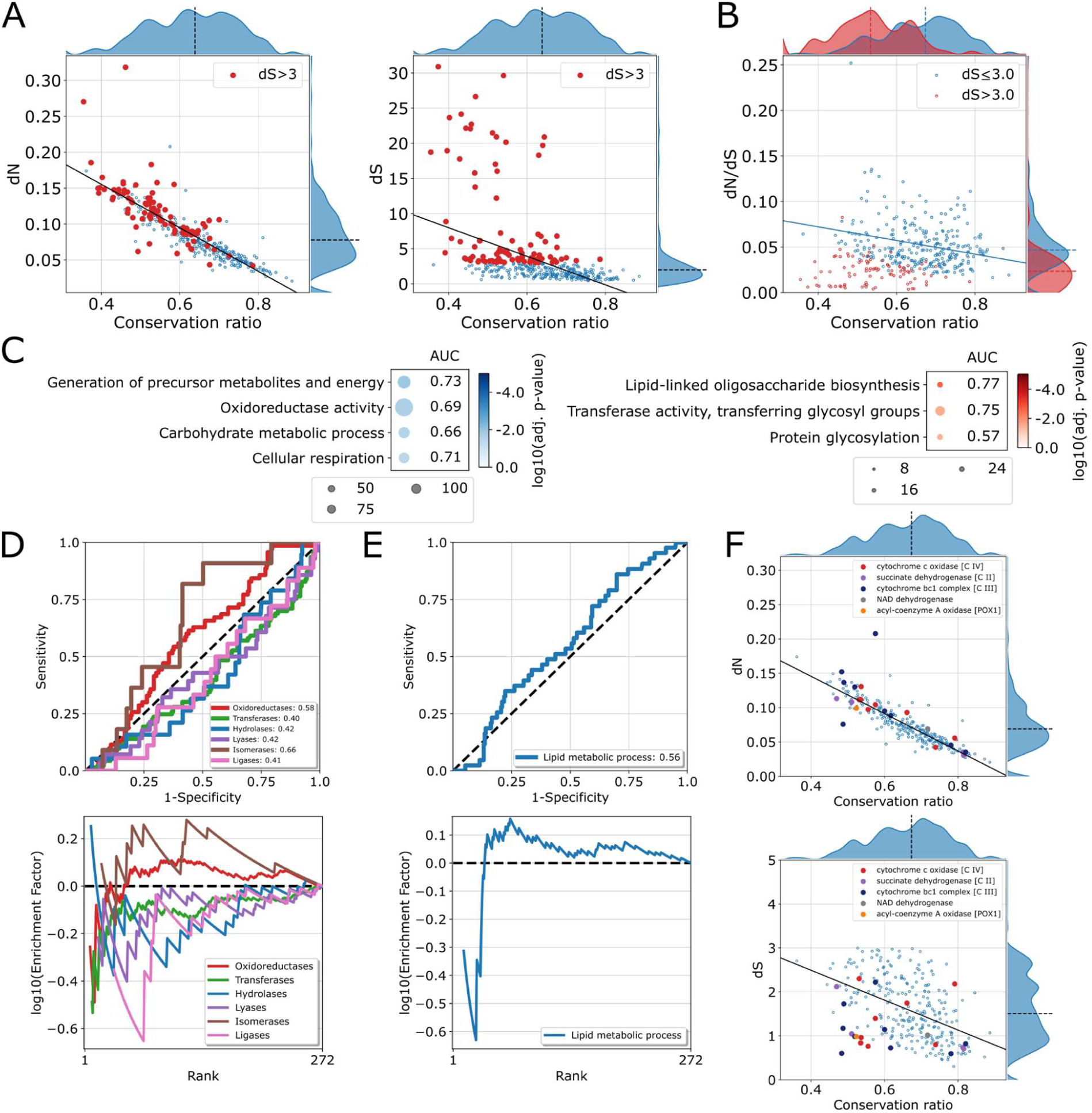
Refinement of the dN/dS calculation to remove skewed values. A) dN (left) and dS(right) versus mean conservation ratio. Red dots denote a dS>3, the solid line indicates the best linear fit and the dashed line denotes the axis median. B) dN/dS versus mean conservation ratio. Colouring according to A). The solid line indicates the best linear fit for the blue highlighted orthogroups and the dashed lines denote the axis median for the two groups. C) Significantly enriched pathways and GO slim terms of the blue (dS≤3) and red (dS>3) highlighted orthogroups with respect to the full set. Size indicates the number of genes associated with the term or pathway in our selection, and colour indicates adjusted p-value. In addition, the area under the receiver operating characteristic (ROC) curve (abbrev. AUC) is shown. D+E) ROC-curve (top) and enrichment factor (bottom) for D) enzyme classes and E) for the GO term “lipid-metabolic process” according to dN/dS. The dashed line indicates random sampling. F) dN (top) and dS (bottom) respectively versus the mean conservation ratio for orthogroups with dS<3. Orthogroups that contain proteins of the electron transport chain as well as Pox1p are highlighted.

**Figure S6.2:**
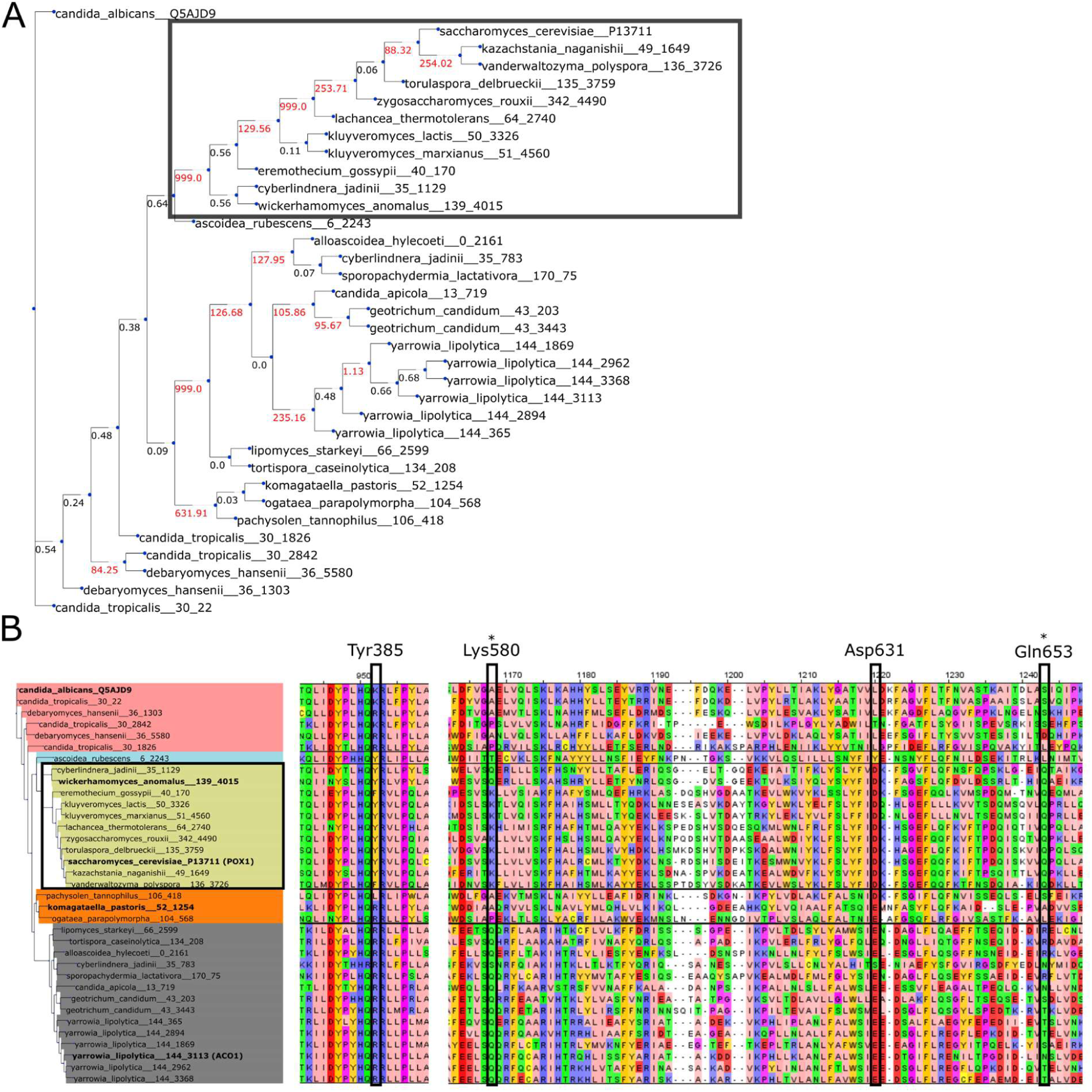
Positive selection in Pox1p. A) The protein tree for the orthogroup containing *S. cerevisiae* Pox1p (OG1122). The red numbers indicate branches for which branch testing (M1 strategy) indicated positive selection dN/dS>1.1, and for which branch site testing was conducted. The dark square indicates the branch containing *S. cerevisiae* and *W. anomalous* orthologs for which residues under positive selection are highlighted in Figure 6C. B) Multiple sequence alignment of selected regions of POX1 orthologs. Asterix indicates residues under positive selection in the branch highlighted with the black box on the left. Other highlighted residues interact with positively selected residues in the structure (See Figure 6C).

**Figure S6.3:**
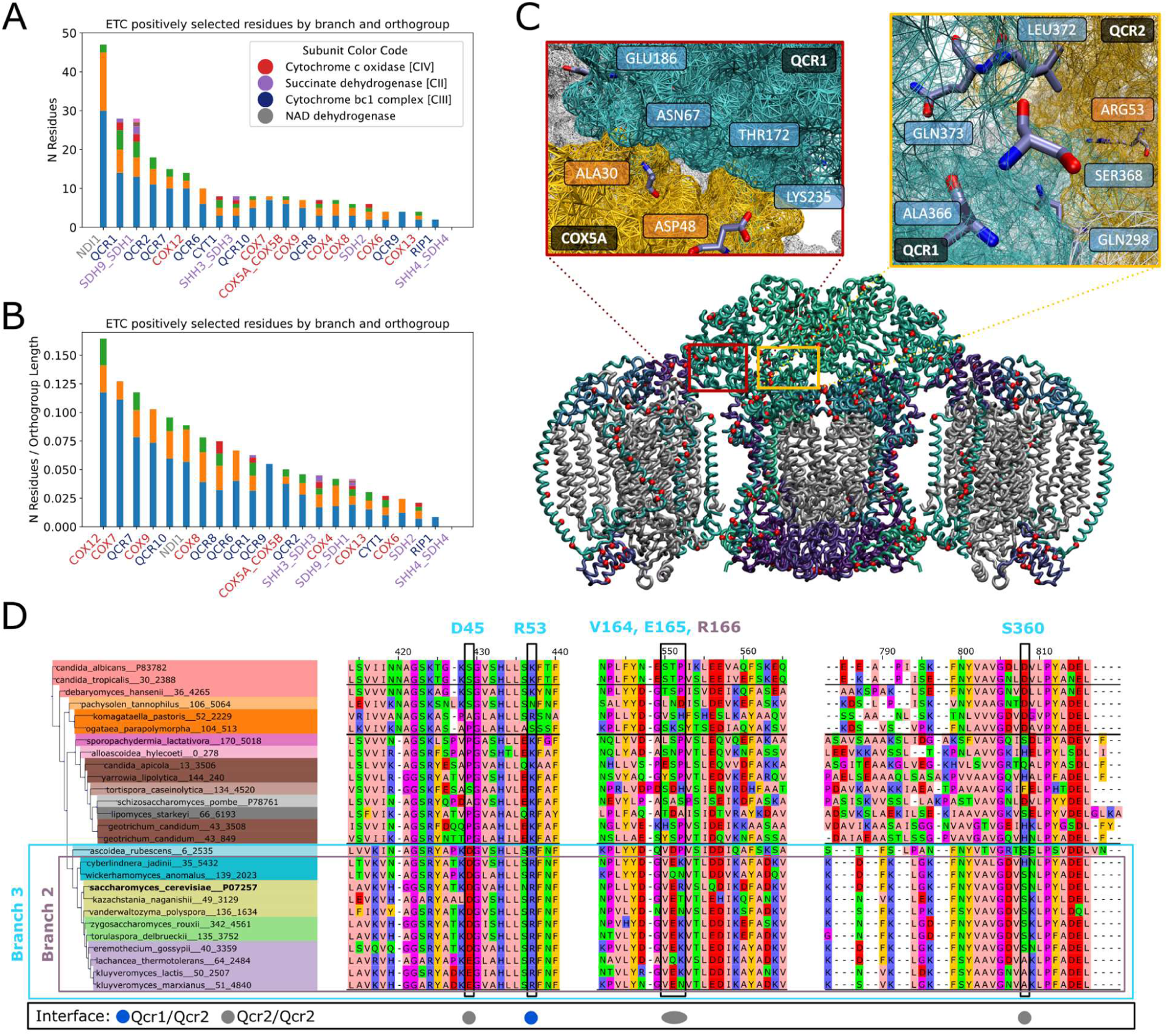
Positive selection in the electron transport chain (ETC). A) Number of positively selected residues for each orthogroup in the ETC. Colours in the bars indicated different branches within the protein tree for which the branch site test for positive selection was conducted. Label colour indicates the ETC subunit of the orthogroup (per legend). B) As in (A) but normalised by mean protein length for each orthogroup. C) Interaction interfaces for Cox5Ap and Qcr1p as well as for Qcr1p and Qcr2p. The respective positions in the complex are denoted in coloured boxes. D) Multiple sequence alignment of Qcr2 highlighting residues detected as under positive selection in designated branches.

